# A molecular framework for proximal secondary vein branching in the *Arabidopsis thaliana* embryo

**DOI:** 10.1101/2021.10.12.464048

**Authors:** Elizabeth Kastanaki, Noel Blanco-Touriñán, Alexis Sarazin, Alessandra Sturchler, Bojan Gujas, Antia Rodriguez-Villalon

## Abstract

- The establishment of a closed vascular network in foliar organs is achieved through the coordinated specification of newly recruited procambial cells, their proliferation and elongation. An important, yet poorly understood component of this process, is secondary vein branching; a mechanism employed in *Arabidopsis thaliana* cotyledons to extend vascular tissues throughout the organ’s surface by secondary vein formation.
- To investigate the underlying molecular mechanism in vein branching, we analyzed at a single-cell level the discontinuous vein network of *cotyledon vascular pattern 2 (cvp2) cvp2-like 1 (cvl1)*. Utilizing live-cell imaging and genetic approaches we uncovered two distinct branching mechanisms during embryogenesis.
- Similar to wild type, distal veins in *cvp2 cvl1* embryos emerged from the bifurcation of cell files contained in the midvein. However, the branching events giving rise to proximal veins are absent in this mutant. Restoration of proximal branching in *cvp2 cvl1* cotyledons could be achieved by increasing *OCTOPUS* dosage as well as by silencing of *RECEPTOR LIKE PROTEIN KINASE 2 (RPK2)* expression. The RPK2-mediated restriction of proximal branching is auxin and CLE-independent.
- Our work defines a genetic network conferring plasticity to *Arabidopsis* embryos to adapt the spatial configuration of vascular tissues to organ growth.

## Introduction

The appearance of a continuous vascular network in plants, as means of water and nutrient exchange among organs, greatly contributed to their conquering of a wide range of terrestrial ecosystems (Lucas et al. 2013; Agusti and Blazquez 2020). The evolution of plants is characterized by the selection of a spatial arrangement of vascular strands (vascular patterns) that maximize their overall functionality (Lucas et al. 2013). In *Arabidopsis thaliana* (Arabidopsis), the vascular pattern of foliar organs is generated and maintained through the formation of continuous procambial cell files; these are further organized in vascular bundles comprising the conductive tissues phloem and xylem (Lavania et al. 2021). While in most species the patterning of leaf vascular tissues exhibits a high degree of plasticity (Scarpella 2017), the robust and reproducible patterns of the vein network in Arabidopsis cotyledons offer an ideal model to identify the positional and molecular cues underlying this process. The vascular network in these organs includes a single primary vein (midvein) that extends along the central part of this organ, and ensures the connection to the stem’s vascular system (Scarpella 2017). A pair of secondary veins diverge from the midvein and extend toward the cotyledon margins as this organ expands laterally due to the proliferation of plate meristematic cells (Fig. 1O) (Scarpella 2017; Tsukaya 2021). These vascular cells are surrounded by mesophyll cells, which have been proposed to limit vein propagation in Arabidopsis leaves through their differentiation, terminating the vein path (Scarpella et al. 2004). Similar to vascular cells, mesophyll cells derive from ground meristem (GM) cells located in the subepidermal layer of cotyledons (Lavania et al. 2021). During the specification of GM cells into either mesophyll or vascular cells, positional cues determine the acquisition of both identities and thus cell types (Lavania et al. 2021). The widely accepted model for cotyledon/leaf vascular cell specification, the auxin canalization model, supports that a directional auxin flow acts as a pre-pattern and reinforces vascular cell identity delineating an incipient vascular path along the cells that have been most exposed to the auxin flow (Scarpella et al. 2006; Lavania et al. 2021). In particular, specific subepidermal GM cells of the cotyledon will acquire pre-procambial identity (Scarpella 2017), which is associated with the expression of auxin-related genes such as *MONOPTEROS (MP)* or *ARABIDOPSIS THALIANA HOMEOBOX 8 (ATHB8)*. The subsequent elongation of pre-procambial cells results in their transition to procambial cells (Lavania et al. 2021).

**Figure 1.**
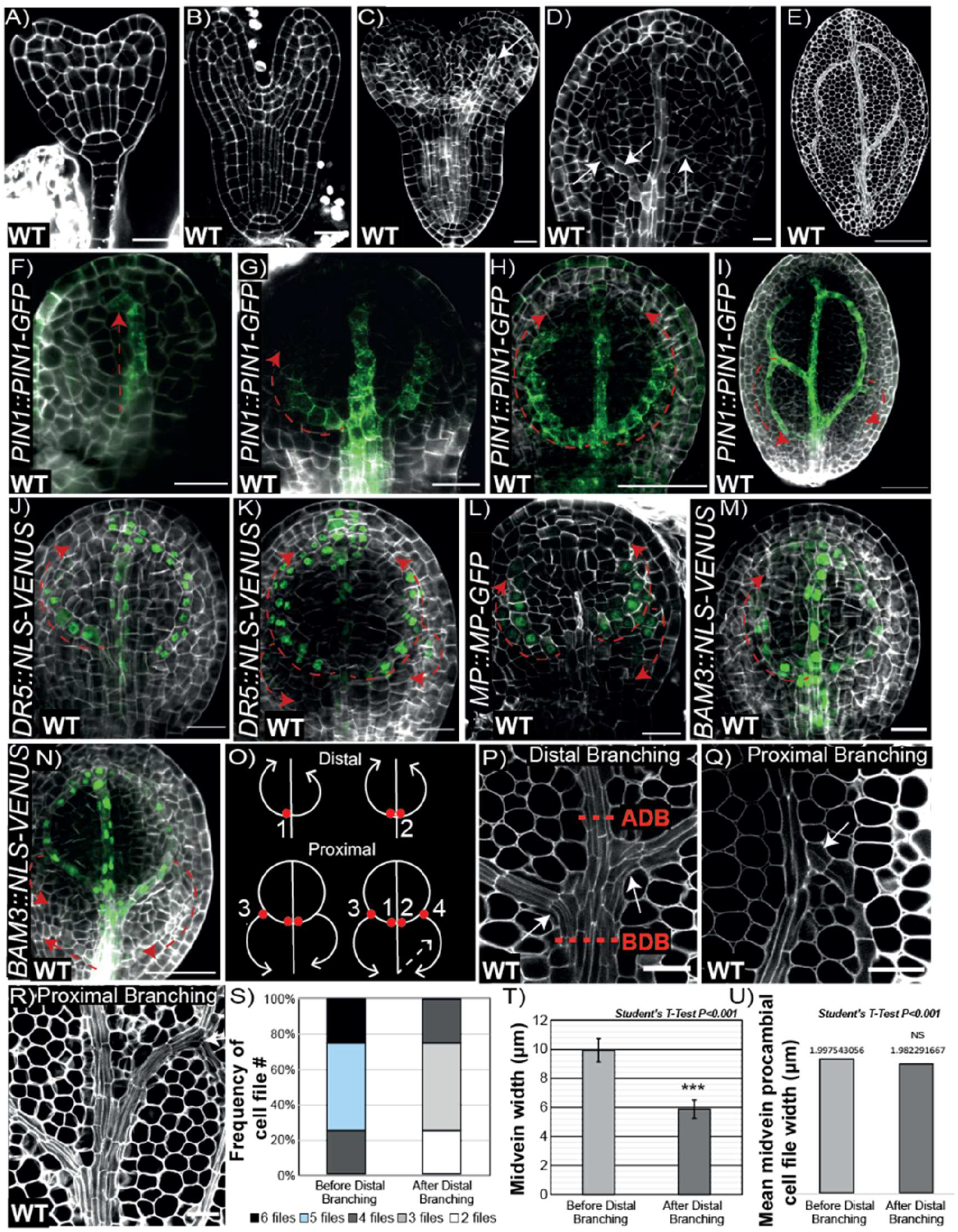
Vein progression and branching in torpedo embryos. **A-B)** Representative pictures of mPS-PI cell wall-stained embryos in late transition stage (A) and late heart stage (B). **C-D)** Analysis by confocal microscopy of extracted embryos from ovules and stained with the cell wall dye SR2200 Renaissance. (C) Embryo in transition between heart and early torpedo in which the future midvein is marked by a white arrow. (D) Cotyledon of early torpedo stage in which the secondary vein formation can be detected. Note that in D, based on the elongated morphology of procambial cells, secondary vein formation is initiated from the midvein and progressing upwards towards the top of the cotyledon (marked by white arrows). **E)** Cotyledon of mature embryo in which the cotyledon vein network is completed. **F-I)** Representative images of *PIN1∷PIN1-GFP* early and late torpedo stage embryos stained with SR2200 as a cell wall counter stain showing the progression of midvein or primary vein (F), distal secondary vein (G-H) and proximal secondary vein formation (I). Dashed red arrows represent the directionality of the forming veins. **J-K)** Cotyledons from early torpedo stage embryos harbouring *DR5∷NLS-VENUS* showing the progression of distal secondary veins (J) as well as the initiation of the proximal secondary veins (K). **L-N)** Early torpedo stage embryos harbouring *MP∷MP-GFP* (L), and *BAM3∷NLS-3XVENUS* (M, N) showing cotyledon proximal vein formation occurring in a tip-to-base manner (n=25/27), except in N, in which proximal veins also proceeds in a base-to-tip manner (n=2/27). **O)** Scheme representing the proposed branching sites of distal and proximal secondary veins. Distal branching points 1 and 2 are represented by the red dots and the direction of vein formation is represented by the white arrows. Proximal secondary vein branching, branching points 3 and 4 (red dots) and the direction of vein formation (white arrows) are represented in the lower panels. Note that distal vs proximal secondary vein formation occurs normally in opposing directions. The rare appearance of proximal veins in base-to-tip manner is represented by a dashed white arrow. **P-R)** Representative images of proximal and distal branching in embryonic cotyledons stained with mPS-PI and visualized by confocal microscopy. ADB: after distal branching; BDB: before distal branching. **S-U)** Quantification of the frequency of appearance of the indicated number of cell files (S), average midvein width (T) and average midvein cell file width (U) in the region marked as ADB and BDB. Note that the difference represented in (T) are not due to differences in the width of procambial cells in these regions, as indicated in (U). Scale bars represent 20μm in A-D, J, L, N and O and 50μm in E, H, I, K, P.

While auxin has been considered the necessary and sufficient signal for vein formation (Lavania et al. 2021), the molecular regulators underpinning the branching of vascular tissues have been poorly explored. Furthermore, the auxin canalization model itself cannot solely explain the appearance of an intermittent vascular network in vein mutants such as *cotyledon vascular pattern 2 (cvp2) cvp2-like 1 (cvl1)* (Carland and Nelson 2009). In particular, the simultaneous depletion of phosphatidylinositol 4,5-bisphosphate [PtdIns(4,5)P_2_] phosphatases CVP2 and its homologous CVL1 results in a reduction in vascular complexity and the appearance of off-path vascular islands (Carland and Nelson 2009). Initially, CVP2 and CVL1 were suggested to generate PtdIns4P, which is required to activate ARF-GAP SCARFACE (SCF) (Naramoto et al. 2009). While the discontinuous vascular network observed in *scf* was related to its role in controlling PIN1 endocytosis (Naramoto et al. 2009), further studies have demonstrated a wider range of subcellular activities such as polarity establishment, which CVP2 and CVL1 modulate. In particular, augmented PtdIns(4,5)P2 levels enhanced subcellular trafficking towards the vacuole, a process that has been associated with the misspecification of root protophloem and companion cells (Gujas et al. 2020; Rodriguez-Villalon et al. 2015). The root vascular phenotypes associated with the skewed ratio of PtdIns4P and PtdIns(4,5)P_2_ can be rescued by introducing *receptor protein like kinase 2 (rpk2)* mutation into a *cvp2 cvl1* background (Gujas et al., 2020). *RPK2* and its homolog *RECEPTOR PROTEIN LIKE KINASE 1(RPK1)* were first described during embryogenesis to control protodermal identity, as evident by the expanded vascular domain found in *rpk1 rpk2* globular embryos (Nodine et al. 2007).

The morphological complexity of vascular tissues has hindered thus far our understanding of the sequence of events resulting in the establishment of the embryonic vein pattern. Here, we revise the directionality of cotyledon vein formation during embryogenesis. We observed that two different mechanisms initiate the branching of distal secondary veins (the secondary veins located in the distal region of the cotyledon) and proximal secondary veins (the secondary veins that emerge from the distal veins in the proximal region of the cotyledon), and that the progression of both types of secondary veins exhibit opposite directionality. While distal branching involves the bifurcation of the cell files comprised in the midvein, proximal secondary veins arise from the branching of distal veins by a poorly understood mechanism. Although distal branching involves PIN1 function, polar auxin transport seems not to control vascular cell fate commitment, as revealed by the continuous vascular network observed in high order *pin* mutants. Furthermore, we report that both CVP2 and CVL1 are necessary to promote proximal branching, which is counteracted by the activity of RPK2. Our work demonstrated that silencing *RPK2* expression partially restores *cvp2 cvl1* cotyledon vein network complexity by promoting vein branching, which is necessary to create proximal secondary veins. Through transcriptomic profiling and genetic assays, we found that a reduced activity of RPK2 restores the impaired cotyledon vein pattern of *cvp2 cvl1* embryos independent of auxin and vascular-specific CLAVATA SURROUNDING EMBRYO (CLE) peptides. By expressing *RPK2* at the cotyledon margins, plants establish a vein network boundary by which veins do not extend towards this area. Additionally we showed that the positive vascular regulator *OCTOPUS (OPS)* promotes proximal branching. Together, our work supports a genetic network by which the positive regulation of vein branching by *CVP2*, *CVL1* and *OPS* is limited by *RPK2*, modulating the vein pattern to organ outgrowth and in turn, maximizing the functionality of vascular tissues.

## Materials and methods

### Plant material and growth conditions

Arabidopsis ecotype Columbia-0 was used as a wild-type control in all cases with the exception of the experiments related to *pin1-*3, *pin1-*5 and and *clv3-7*, in which Ler was used as a control, respectively. Seeds of *pin1-3*, pin1-5, *pin1,3,6,4,7,8* and *PIN1∷PIN1-GFP* were kindly provided by Dr. Friml (Institute of Science and Technology, Austria), Dr. Miguel Perez-Amador (Institut of Plant Molecular Biology of Valencia, Spain) and Dr. Scarpella (University of Alberta) whereas seeds of *aux1-21 lax1* were provided by Dr. Fankhauser (University of Lausanne). *cle45.cr2* and *clv3* loss-of-function mutants were provided by Dr. Takashi Ishida (Kumamoto Health Science University, Japan) and Dr. Hamant (ENS Lyon), respectively. The transgenic lines *SHR∷SHR-GFP* and *MP∷MP-GFP were* provided by Dr. Vermeer and Dr. Weijers, respectively. *CVP2∷NLS-3xVENUS*, *CVL1∷NLS-3xVENUS, CVP2∷GUS and CVL1∷GUS* have been previously described (Rodriguez-Villalon et al. 2014; Rodriguez-Villalon et al. 2015; Gujas et al. 2020; Carland and Nelson 2009). Likewise, *rpk2, cvp2 cvl1*, *amiRPK2 cvp2 cvl1* and *ops* were reported elsewhere (Gujas et al. 2020). *MP∷MP-GFP, SHR∷SHR-GFP, DR5∷NLS-VENUS, PIN1∷PIN1∷GFP, AUX1∷AUX1-YFP* and *OPS∷OPS-GFP* translational fusion reporter lines in distinct genotypes were obtained by crossing these lines with the indicated loss-of-function mutants or have been previously published (Rodriguez-Villalon et al. 2015). Seeds were surface-sterilized, stratified at 4°C and grown vertically on 0.5x MS plates under standard continuous-light growth conditions. Seedlings were transferred to soil and grown in 5×5 cm pots in peat-based compost medium in a walk-in chamber at constant 23°C, 65% humidity in a 16h photoperiod and light intensity of 250μmol photons m^−2^ sec-^1^ until flowering when siliques were collected to extract embryos.

### Cloning and plant transformation

All constructs were generated using double or triple Multi-Site Gateway system following the handbook instructions. To clone *ABI3* promoter, 2kb genomic DNA region was PCR-amplified and introduced via pDNRP4-P1r (Invitrogen) to generate pENTRY-ABI3. CLV3 and CLE45 were amplified using the following primers CLV3_XmaI_F “TAACCCGGGATGGATTCGAAGAGTTTTCTGC” CLV3_SpeI_R “CGCCACTAGTTCAAGGGAGCTGAAAGTTGT” CLE45_XmaI “TAACCCGGGATGTTGGGTTCCAGTACAAGA” CLE45_SpeI_R CGCCACTAGTTTAAGAAAATGGCTGAGCTTTGT” and the PCR products were cloned into pENTRY vectors by standard procedures and further recombine together with pENTRY-ABI3 into the destination vector pEDO 097. *BAM3* promoter (2139 bp) was amplified using the following primers: pBAM3_attB4_F: GGGG ACA ACT TTG TAT AGA AAA GTT GCC CTGCTTCCCTAGTTTATCTAATAAATCTGATG and pBAM3_attB1r_R: GGGG AC TGC TTT TTT GTA CAA ACT TGG TGTAACATCAGAAAAATAAAAACAAAAATTTGTCC and fused to *NLS-3xVENUS* construct as previously described (Gujas et al., 2020). Transgenic plants were generated using floral-dip transformation techniques as previously described (Gujas et al. 2017).

### Confocal microscopy

Mature embryos were dissected from seed coats and stained utilizing the Modified Pseudo Shiff Propidium Iodide (mPS-PI) method according to (Truernit et al. 2008) and then imaged using a ZEISS LSM 780 confocal microscope. Embryos in globular stage and onwards were visualized after being dissected from siliques and fixed in 75% ethanol and 25% acetic anhydride for 24hours at 4 degrees Celsius. After the fixing step, embryos were washed, incubated in 1% period acid and rinsed with water following incubation in Schiff’s reagent and Propidium Iodide. Finally, embryos were mounted on glass slides in chloral hydrated (Sigma-Aldrich 302-17-0). Embryos with fluorescent reporters were imaged using Renaissance staining SR2200 as previously described (Smit et al. 2020). Pictures showing the localization of protein in the roots were obtained using a two-photon laser Leica SP8 microscope. 6-day-old roots were stained for 5 min in a 10 μg/ml aqueous solution of propidium. For esthetical reasons, images were rotated and displayed on a matching background. All image processing was performed using ImageJ software. Procambial cell file width was measured on individual procambial cells belonging to the midvein using ImageJ and represented as a mean before and after distal branching. Midvein width was measured as the total width of all procambial cell files comprising the midvein before and after distal branching. Intensity ratios to assess polar protein distribution in procambial and protophloem cells were quantified using ImageJ by measuring mean intensity for each region of interest (ROI). Basal, apical and lateral membranes’ signals were normalized by the total ROI area. The mean of all cells quantified is represented as the intensity ratio of each protein in the corresponding genotype.

### mRNA sequencing of Arabidopsis embryos and data analysis

50 torpedo staged embryos of each genetic background, WT, *cvp2 cvl1* and *pRPK2∷amiRPK2 cvp2 cvl1* were manually dissected out of the ovule with needles and immediately placed in TRIzol reagent (Ambion) and grounded with a sterile pestle. Two or three biological replicates (50 embryos) were collected and analyzed. All samples were frozen in TRIzol and kept at −80°C until RNA extraction. RNA extraction was completed by incubating samples at 60°C for 30 minutes and then purified according to (Palovaara et al. 2017). The resulting RNA was cleaned up and concentrated using the RNeasy MinElute Cleanup Kit Qiagen (74204) according to (Palovaara et al. 2017). Samples were eluted with 14uL RNase-Free water and stored at −80°C. Isolated RNA displayed a RNA Integrity Number (RIN) ranging from 6.8 to 7.7. mRNA-seq libraries were generated with the Smart-seq2 kit (Agilent) and subsequently sequenced in an Illumina NovaSeq 6000 in the Functional Genomics Center Zurich (FGCZ). Data have been stored on the Gene Expression Omnibus (GEO) with accession number GSE178241. Quality validation was carried on using FastQC (https://www.bioinformatics.babraham.ac.uk/projects/fastqc/), after which adapter sequences were removed using Trimmomatic (v0.39, PMID: 24695404) with options SE -phred33 ILLUMINACLIP:trimmed/NexteraPE-PE.fa:2:30:10 SLIDINGWINDOW:4:15 MINLEN:50. Trimmed reads were then aligned onto Arabidopsis thaliana TAIR10 Ensembl genome and genes annotation (retrieved from igenome) using HISAT2 (v2.2.1; PMID: 31375807) with options -k 10 --max-intronlen 1000--known-splicesite-infileTAIR10_splicesites.txt after applying hisat2_extract_splice_sites to TAIR10 Ensembl genes annotation. Reads count table for annotated genes was generated using the featureCounts function (v2.0.1) from the Subread package (PMID: 24227677) with options -O -M -T 10—largest Overlap-- minOverlap 10 --primary. Differential analysis was performed using DEseq2 (v1.22.2, PMID: 25516281). Genes with a p-adjusted value (padj) lower than 0.05 were considered as differentially expressed.

### Quantitative reverse transcription-PCR (RT-qPCR)

Total RNA was isolated with a RNeasy Plant Mini Kit (QIAGEN) according to the manufacturer’s instructions. cDNA was prepared from 2μg of total RNA with Thermoscientific RevertAID First Strand cDNA Synthesis Kit following manufacturer’s instructions. Resulting cDNA was diluted 1:10 in ddH_2_O and 2 μL of the resulting dilution were used in the PCR reaction. qPCR was prepared using KAPA SYBR FAST qPCR mix. All reactions were performed in triplicates and expression levels were normalized to those of *PDF2*. Primers sequences are shown below: *PDF2*_Fw: 5’-TAACGTGGCCAAAATGATGC-3’; *PDF2*_Rv: 5’-GTTCTCCACAACCGCTTGGT-3’ (Czechowski et al. 2005); *AUX1*_Fw: 5’-GGATGGGCTAGTGTAAC-3’; *AUX1*_Rv: 5’-TGACTCGATCTCTCAAAG-3’ (Dindas et al. 2018); *PIN1*_Fw: 5’-ACAAAACGACGCAGGCTAAG-3’, *PIN1*_Rv: 5’-AGCTGGCATTTCAATGTTCC-3’ (Heisler et al. 2005); *OPS*_Fw: 5’-GACAGGTCTAGTAGCTCCATGAGG-3’; *OPS*_Rv: 5’-AGCTTTGGCTCGTCCATATCCG-3’. *OPS* primers were designed using QuantPrime.

### Histology and GUS staining

Developing cotyledons were imaged at 7 or 8 days as indicated. Cotyledons and leaves were fixed with 3:1 ethanol: acetic acid, dehydrated in 80% ethanol, and then 100% treated with 10% sodium hydroxide for 1hour at 37°C and mounted in 50% glycerol. Black and white images were taken and brightness and contrast were adjusted using ImageJ. To visualize GUS staining, we used a staining buffer as previously described in (Carland and Nelson 2004) with 2mM 5-bromo-4-chloro-3indol-b-D-glucuronide and 1mM potassium ferricyanide and ferrocyanide. We considered branching points (BP) as lateral/secondary veins bifurcated from both midvein and distal secondary veins (see model shown in Fig.1O).

### Seedling treatments

To perform NPA and CLE treatments, 4-day-old seedlings grown in MS were transferred to a media supplemented with or without NPA (10μM), CLV3 (5nM), CLE45 (20nM), CLE25 (100nM) and CLE26 (150nM) for 5 days. The quantification of the root sensitivity to NPA treatment was performed by measuring the length of the primary root using ImageJ.

## Results

### Two distinct branching mechanisms control cotyledon vein network formation in torpedo stage embryos

To gain further insight into the contribution of procambial cell identity acquisition and cell proliferation in cotyledon vein network complexity, we first sought to characterize the cotyledon vascular ontogeny during embryogenesis. The emergence of procambial cell files can be detected by confocal microscopy based on their characteristic narrow and elongated morphology. Confocal microscopy analysis of Renaissance stained embryos revealed the appearance of procambial cell files at early torpedo stage, when a midvein could be detected within the cotyledons (Fig. 1A-C). While the complete establishment of the vein network was detectable in late torpedo stage, elongated narrow cells diverging from the midvein could be observed at earlier developmental time points (what we termed intermediate torpedo stage) (Fig. 1D, E). To further corroborate these observations, we monitored the auxin efflux PIN1 protein, whose distribution in torpedo embryos is restricted to procambial and protodermal cells (Fig. 1F). We observed a progressive formation of the midvein concomitant with the base-to-tip appearance of distal secondary veins (Fig. 1F-I, O). The morphological progressive directionality of the distal secondary veins was corroborated through monitoring auxin response by means of *DR5∷NLS-VENUS* and by the gene expression analysis of *MP* (pre/procambial) as well as the root phloem regulator *BARELY ANY MERISTEM 3 (BAM3)* (Fig. 1J-N,O) (Scarpella et al. 2004; Przemeck et al. 1996; Rodriguez-Villalon et al. 2014). Contrary to the base-to-tip growth of the distal secondary veins, PIN1 tagged with a green fluorescence protein (GFP) polarly accumulates at the basal membrane of the vein network procambial cells as previously reported (Fig. 1G,H) (Scarpella et al. 2006). Since the distal vascular strand is extended in a base-to-tip manner, the polar accumulation of PIN1 at the basal membrane appears to act as a reinforcement identity mechanism in cells already committed to vascular cell fate. To assess the origin of the procambial cell files resulting in distal secondary vein formation we analysed the region near the base of the cotyledon. Here we observed that the number of cell files that constitutes the midvein is higher before distal secondary vein formation *vs* right after (Fig. 1P-S). In contrast, the number of cell files comprising the proximal and distal veins appears to be similar (Fig. 1P-U). These observations indicate that while distal secondary veins directly diverge from the cell files comprised in the midvein, the branching of proximal veins appears to be different and follow a yet-to-be described mechanism.

### In *cvp2 cvl1* mutants GM cells fail to commit to procambial cell identity

To better understand the spatio-temporal arrangement of cotyledon vascular formation during embryogenesis, we decided to exploit the discontinuous *cvp2 cvl1* cotyledon venation pattern as a model. Consistent with previous studies (Carland and Nelson 2009), 7-day old *cvp2 cvl1* cotyledons exhibit a discontinuous and more simplified cotyledon vein network (Fig. 2A-C) (Carland and Nelson 2009). Yet, the midvein was always morphologically intact based on cotyledon clearing visualization techniques (Fig. 2B). These vascular defects originated during embryogenesis, as manifested by the analysis of cotyledons in mature embryos stained by pseudo-schiff-propidium iodide (mPS-PI) (Fig. 2D-G). At this developmental stage, procambial cells can be observed as elongated, narrow elements while surrounding mesophyll cells are rather spherical (Fig. 2E, G). The latter morphology can be detected in between vascular islands, indicating that GM cells in *cvp2 cvl1* fail to commit to procambial cell identity (Fig. 2G) and become mesophyll cells instead. This notion was corroborated by the lack of *ATHB8* expression in the spherical cells flanking *cvp2 cvl1* vascular islands, whereas a continuous expression of this gene within the cotyledon vein network can be detected in wild type embryos (Fig. 2H-K). Since *CVP2* and *CVL1* are expressed in the vein domain from globular stage onwards (Supporting Information Fig. S2, Fig. 2L-P), we decided next to evaluate the vascular identity domains in early embryonic stages in *cvp2 cvl1*. In globular embryos, the expression of *SHORT ROOT (SHR)* or *MP* marks the onset of vascular formation and delineates the separation of the future ground tissues (De Rybel et al. 2016; Scarpella 2017). At this particular stage, none of the vascular markers analyzed showed an aberrant expression domain in *cvp2 cvl1* embryos with the exception of *MP* in globular embryos, whose expression appears very weak (Supporting Information Fig. S2F-M). Together, these results indicate that the cotyledon vein network discontinuities in *cvp2 cvl1* are not due to impaired vascular identity domains during the early stages of embryogenesis and must occur in later embryonic developmental stages.

**Figure 2.**
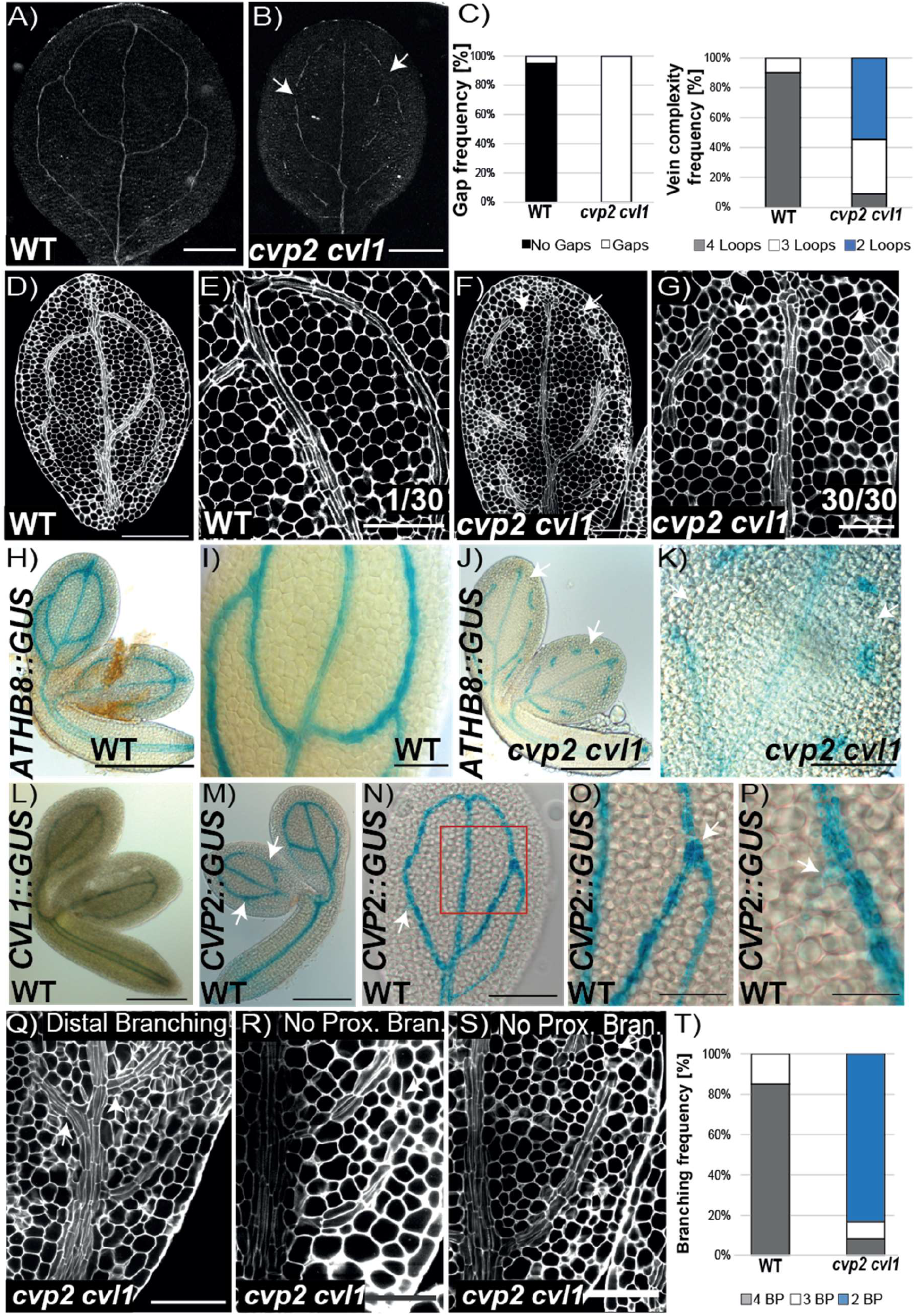
Cotyledon vein defects of *cvp2 cvl1*. **A, B**) Cleared 7-day-old cotyledons of WT and *cvp2 cvl1* imaged with a stereomicroscope in bright field on a black background. **C)** Quantification of the frequency of ground meristem cells surrounding a disconnected vascular islands (gap frequency) and vein complexity in WT and *cvp2 cvl1* cotyledons. n= 20-40 for each genotype. This quantification is part of a bigger experiment which is fully represented in Figure 4G. **D-G)** Confocal microscopy analysis of the vein pattern of mature embryonic cotyledons (vein network at its complete stage) of WT and *cvp2 cvl1* having undergone mPS-PI staining. E and G are magnifications of D and F. White arrows indicate vein gaps. Number of embryonic cotyledons exhibiting gaps is indicated in E and G. **H-K)***ATHB8∷GUS* expression in WT and *cvp2 cvl1* mature embryonic cotyledons. (I, K) Magnifications of the veins displayed in H and J respectively. **L-P)** Expression pattern of *CVL1∷GUS* and *CVP2∷GUS* in embryos. Magnification of an embryonic cotyledon (N) displaying *CVP2* expression in the proximal branching points is shown in (O,P). **Q-S)** mPSI-PI staining of mature embryonic cotyledons of WT and *cvp2 cvl1* showing distal vs proximal branching. Note the two phenotypes observed in *cvp2 cvl1*, when there is a third branching event (R) or not (S). **T)** Quantification of the frequency of the reduced vein complexity observed in each genotype. n= 30. BP: branching points, counted as the initiation (even if not completed) of a new secondary vein. Scale bars represent 50μm in D, F, L, O, P 100 μm in N, and 1 inch in H-K.

### CVP2 and CVL1 activities are required to modulate cell division and proximal branching

Matching the embryonic stages relevant to their expression, we observed a plethora of aberrant morphologies in *cvp2 cvl1* double mutant embryos (Supporting Information Fig. S3). While some embryos resembled wild type (phenotype A), several globular embryos exhibited an excessive number of divisions in the suspensor and hypophysis (phenotype B, Supporting Information Fig. S3). Moreover, cells in the globular embryo of *cvp2 cvl1* display aberrations regarding the number and the orientation of their divisions (Supporting Information Fig. S3). We observed a high embryo abortion rate in *cvp2 cvl1* siliques (ca. 50%), which we believe is partially contributed by aberrant divisions observed mainly in the hypophysis, since an abnormal suspensor development has been reported to be lethal (ten Hove et al. 2015). Although subsequent developmental stages appear morphologically similar to wild type, we cannot exclude the appearance of mild division defects occurring within the vascular domain due to the technical limitations imposed by working with torpedo stage embryos (Supporting Information Fig. S3). However, we observed that proximal branching was absent in the cotyledon vein network of *cvp2 cvl1* (Fig. 2Q-T) even if *CVP2* expression can be detected at the BPs (Fig. 2N-P). In the event of secondary vein formation within the proximal cotyledon region, these procambial cell files appeared to be connected to the basal end of the midvein but without a clear apical branching site (Fig. 2R, S). Indeed, these incipient cell files propagate in a base-to-tip manner (Fig. 2S). Taken together, the presence of an intact midvein and the lack of proximal secondary vein branching imply that the activities of CVP2 and CVL1 are required to initiate proximal secondary veins at the proximal branching point.

### PIN1 polar distribution is not altered in *cvp2 cvl1*

To gain further insight into the vein network branching defects observed in *cvp2 cvl1* embryonic cotyledons, we decided first to assess the role of auxin and its PIN1-mediated transport. Confocal microscopy analysis of PIN1-GFP in *cvp2 cvl1* torpedo embryos showed a normal basal polarization of the auxin carrier in the distal secondary veins (Fig. 3A-D’), which we confirmed by the quantification of PIN1 accumulation in the basal membrane in comparison to a lateral membrane (Fig. 3E) Similarly, analysis of *PIN1-GFP* in root protophloem cells, where PIN1 is polarly accumulated at the basal membrane, did not show any abnormal distribution of this auxin efflux carrier (Fig. 3F-J’). Previous reports have suggested that PIN1 activation requires the function of PIP5K1 and PIP5K2 (Marhava et al. 2020), which catalyze the inverse enzymatic reaction as that of CVP2 and CVL1 (Gujas and Rodriguez-Villalon 2016). To determine the extent to which the vascular phenotypes observed in *cvp2 cvl1* cotyledons may be due to a perturbed PIN1 activity, we decided to analyze the vascular phenotypes of *pin1* seedlings. A plethora of cotyledon defects have been described in distinct *pin1* mutants, including fused cotyledons and aberrant morphologies (Friml et al. 2003). To overcome the impact of defective organogenesis in the analysis of vascular patterning, we decided to focus on separated cotyledons from the loss-of-function *pin1-3* and *pin1-5* mutants. We consistently observed an increased distal branching in *pin1* single mutants while the continuity of the vascular strands appeared intact (Fig. 4A-C, G-I). Likewise, the simultaneous depletion of other vascular PIN carriers such as *PIN3 PIN4 PIN6 PIN7* and *PIN8* (*pin1,3,4,6,7,8*) did not result in a discontinuous vein pattern (Fig. 4E, H, I), although the duplication of the midvein and subsequent bifurcation in distal veins could be found in these mutants (Fig. 4E). These observations indicated that the polar auxin transport represses distal branching. An increase in PIN1 dosage by introducing *PIN1∷PIN1-GFP* (Yanagisawa et al., 2021) (Supplemental Fig. S1) in *cvp2 cvl1* did not aggravate the vascular phenotype of the double mutant (Fig. 3K-N), consistent with our findings showing that *cvp2 cvl1* exhibits defects in proximal branching yet is able to form distal secondary vein branching points (Fig. 2Q, S). To identify the origin of vascular identity failure observed in *cvp2 cvl1* embryonic cotyledons, we decided to analyze auxin response. The deficient activity of the auxin receptors TRANSPORT INHIBITOR RESPONSE 1/AUXIN SIGNALING F-BOX (TIR1/AFB) suppress the formation of secondary veins whereas the midvein appears still intact (Fig.4 F), a phenotype consistent with previous publications (Mazur et al. 2020). To further investigate whether the inability of *cvp2 cvl1* cells to perceive auxin could explain the appearance of a discontinuous vascular network in this mutant, we decided to monitor the distribution of *DR5∷NLS-VENUS*, a widely used auxin response biosensor. Confocal microscopy analysis of embryonic cotyledons showed a positive correlation among cells exhibiting vascular morphology and *DR5* expression (Fig. 4J-M). However, no signal was detected in the cells flanking vascular island (Fig. 4L-M). Taken together, our observations suggest that while auxin cues may be necessary to establish continuous secondary veins in *cvp2 cvl1* embryos, the polar transport of this hormone contributes to regulate the distal branching of secondary veins but not their continuity.

**Figure 3.**
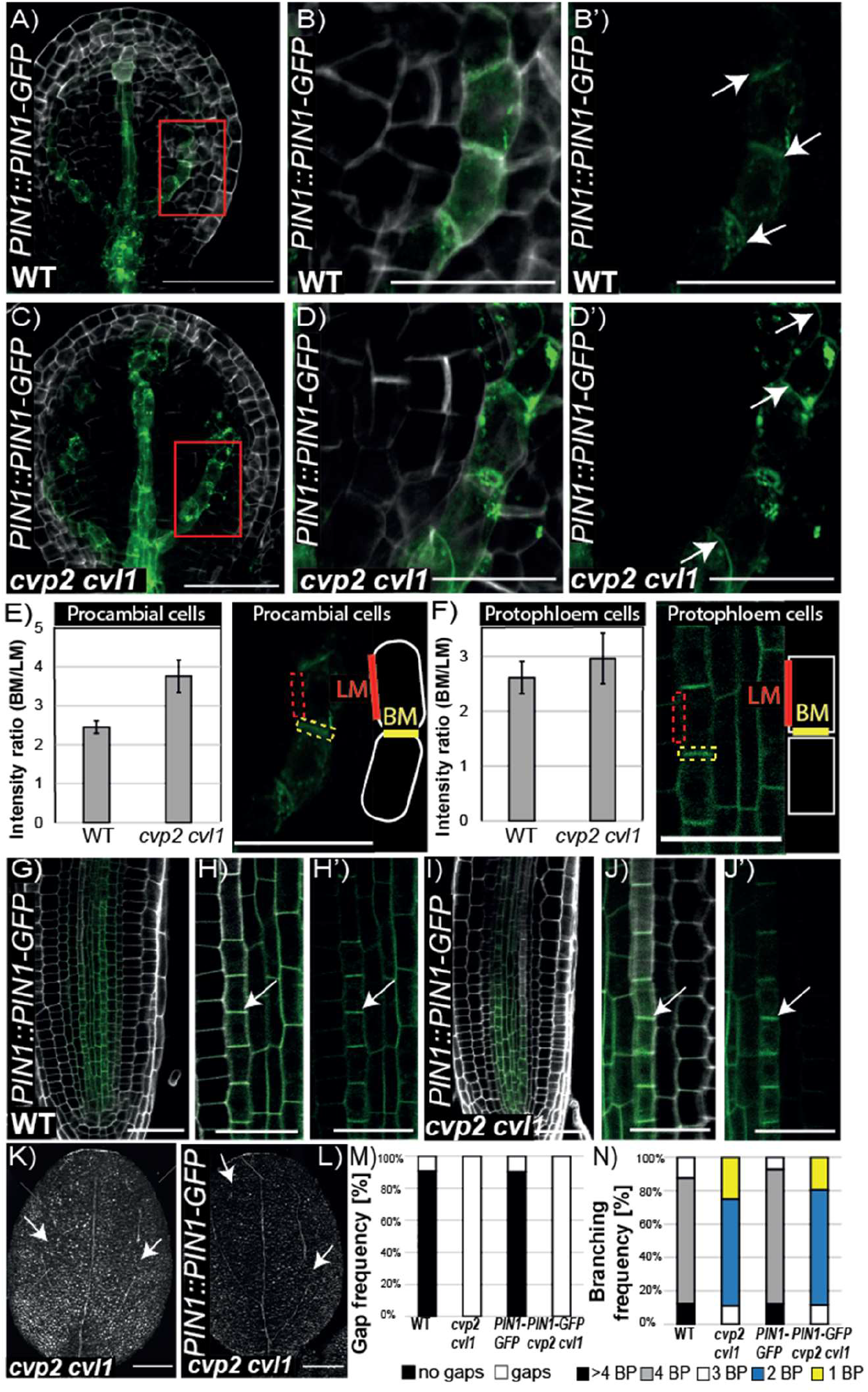
PIN1 polar distribution is not affected in *cvp2 cvl1*. **A-D’)** Confocal microscopy analysis of early torpedo stage embryos of the indicated genotypes stained with SR2200 Renaissance showing PIN1-GFP distribution in distal secondary veins as they are progressively forming upwards. Magnifications of the region squared in red in A) (B, B’) and, in C) (D, D’) are displayed. **E-F)** Quantification of the polar distribution of PIN1-GFP in procambial and protophloem cells as ratio of GFP signal detected in the basal membrane (BM) versus the lateral membrane (LM). A polarity index bigger than 1 is considered a polar distribution. **G-J’)** 6-day-old roots harbouring *PIN1∷PIN1-GFP* in WT and *cvp2 cvl1* background showing PIN1 localization and strong basal polarization in the protophloem strand. Magnification of protophloem differentiating cells in WT (G, H’) and *cvp2 cvl1* (I, J’). Scale bars represent 50μm (A-D’), 20μm (G-J’) and 200 μm (K-L).

**Figure 4.**
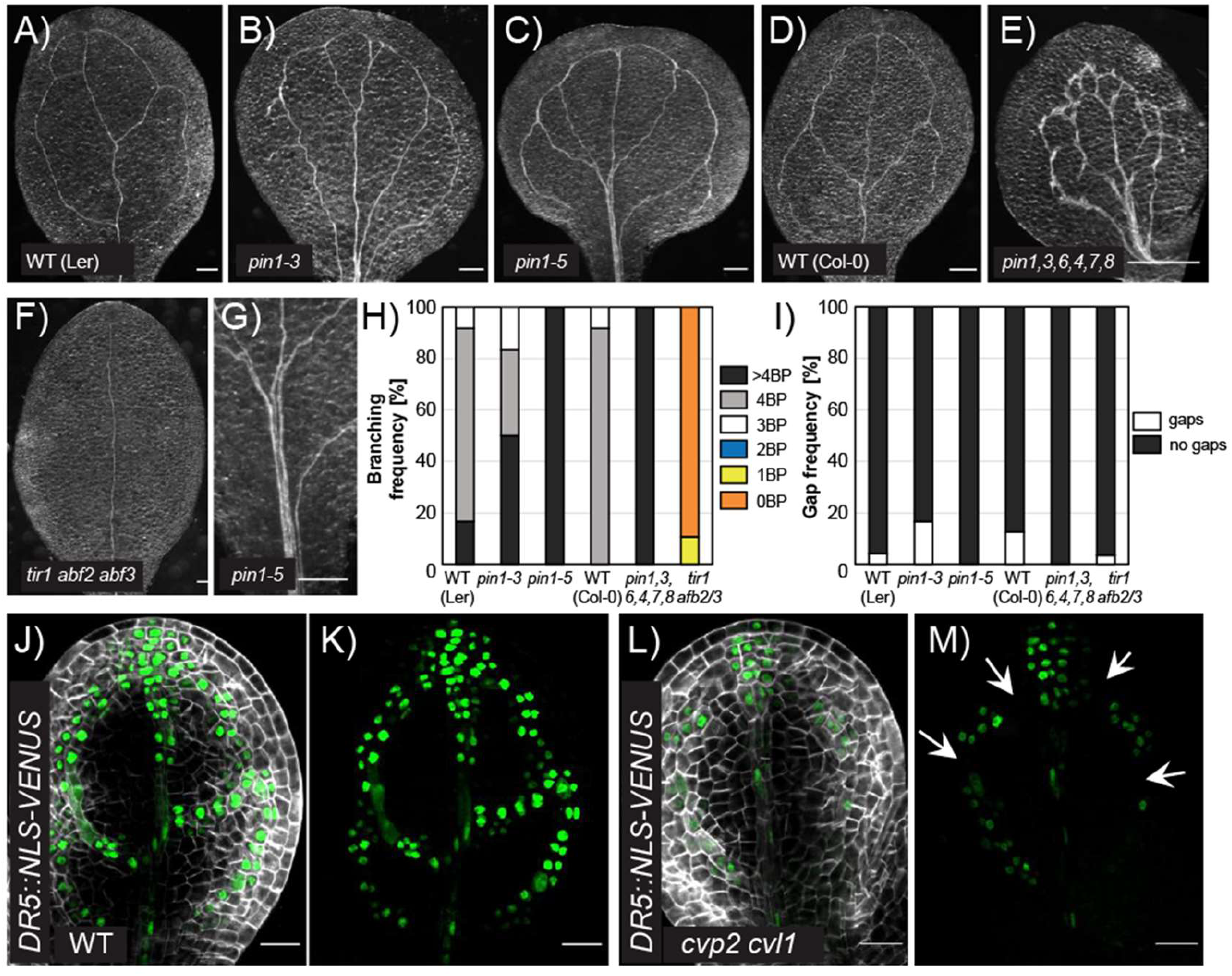
PIN-mediated auxin transport is not involved in modulating proximal branching in embryonic cotyledons. **A-G)** Representative images of 7-day-old cleared cotyledons of the indicated genotypes. Note that *pin1* single mutants are in a Ler background. Magnification of the midvein region where distal branching occurs in *pin1*-5 is displayed in G. **H, I)** Quantification of branching (H) and gap (vascular discontinuities) (I) frequency in the indicated genotypes. n= 30-52 for each genotype. BP: branching points, counted as the initiation (even if not completed) of a new secondary vein. **J-M)** Auxin distribution analyzed by *DR5* expression in WT (J, K) and *cvp2 cvl1* (L, M) embryonic cotyledons counterstained with SR2200 Renaissance. K and M displayed GFP signal. Scale bars in J-M represent 20 μm and in A-F 200 μm.

### Silencing of *RPK2* expression rescues the proximal branching defects of *cvp2 cvl1* embryonic cotyledons

Considering that auxin transport itself cannot solely explain the vascular defects observed in *cvp2 cvl1*, we decided to explore cotyledon vascular ontogeny in plants with a deficient activity of RPK2. We have previously reported that silencing of *RPK2* expression in *cvp2 cvl1* (*amiRPK2*) restores the continuity of the root protophloem strands in this mutant (Gujas et al. 2020). Notably, a partial silencing of *RPK2* expression did not rescue the discontinuous secondary veins of *cvp2 cvl1* (Fig. 5 A, B, D, E). Instead, an increased number of branching points could be observed in these lines (Fig. 5A, B, D-G, K, K’). In light of these results, we decided to further elucidate the potential role of RPK2 in cotyledon vein patterning. Examination of *rpk2-2* vascular pattern revealed the occasional appearance of additional proximal branching points within the embryonic cotyledon vein network, even if a complete/closed additional aerole was rarely observed (Fig. 5C, F, G). Furthermore, divisions giving rise to a bifucarted vein path could also be occasionally detected in mPS-PI stained *rpk2-2* embryos (Fig. 5H-K’). Consistent with previous studies, *RPK2* expression can be mainly detected in the protoderm and in some cells of its adjacent cell file (Fig. 5L, L’) (Nodine et al. 2007). Confocal microscopy analysis of *RPK2* expression in *cvp2 cvl1* however, revealed a slightly broader expression pattern towards the inner cell layers, with is greatest expression at the tip of the cotyledon (Fig. 5M, M’). By repressing the branching of proximal secondary veins, *RPK2* modulates vein complexity while preventing the extension of procambial cell files into the cotyledon margin area.

**Figure 5.**
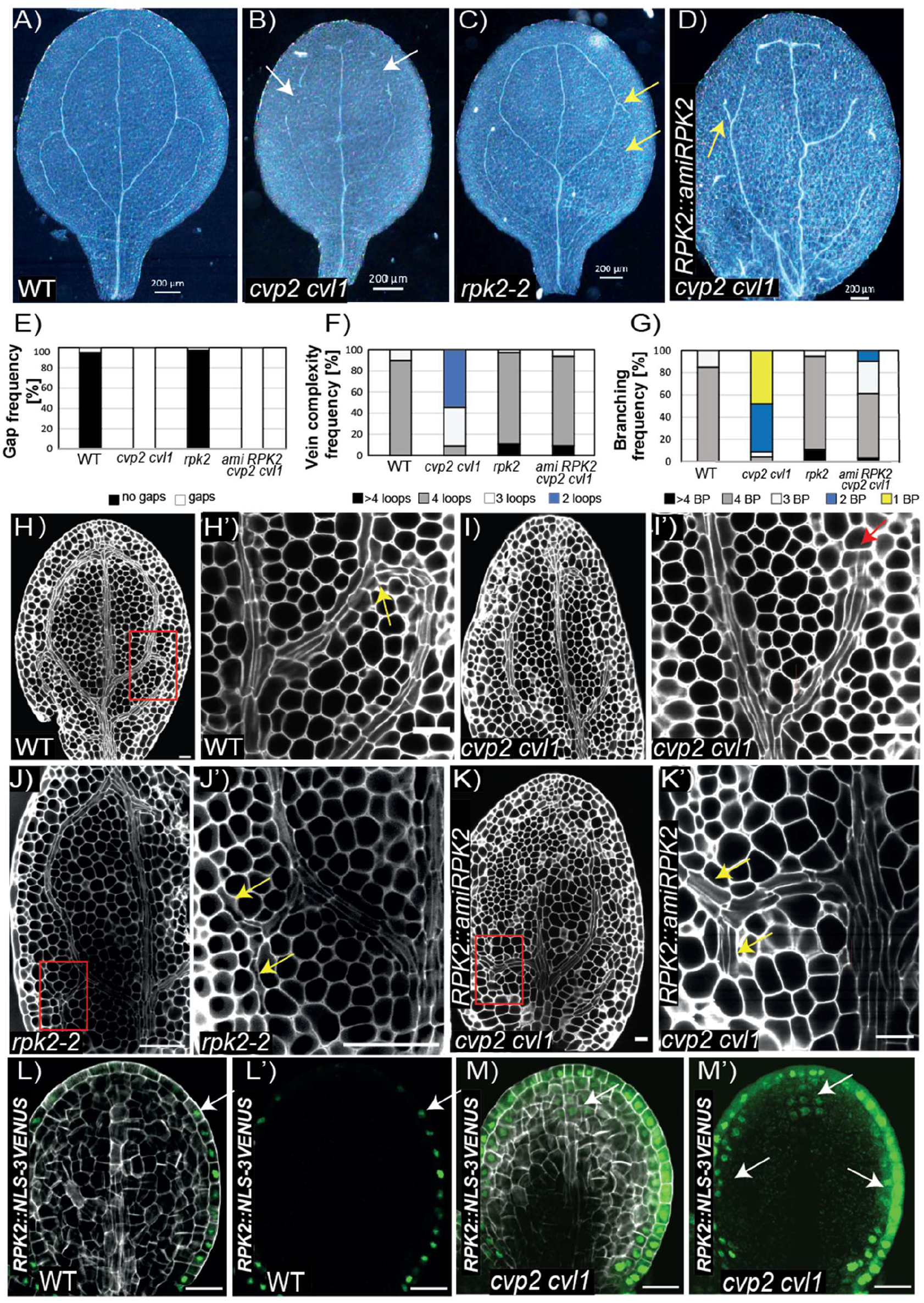
Silencing of *RPK2* expression rescues the branching defects of *cvp2 cvl1*. **A-D)** Analysis of the continuity and complexity of cotyledon vein network in 7-day-old seedlings of the indicated genetic backgrounds. **E-G)** Quantification of gap (E), vein complexity (F) and branching (G) frequency observed in the cotyledons of the plants depicted in A-D. n= 23-50 for each genotype. **H-K’)** mPS-PI stained embryos displaying the vein pattern of the indicated genotypes. H’, I’, J’ and K’ represent a magnification of the squared region represented in H, I, J and K respectively. Yellows arrows mark proximal branching while the red arrow marks the lack of proximal branching. **L-M’)** Confocal microscopy analysis of *RPK2* expression in the cotyledons of torpedo embryos of the indicated genotypes stained with Renaissance. L’ and M’ show only GFP signal. Scale bars represent 200 μm in A-D, 20 μm in H-M’.

### The partial *RPK2-*mediated restoration of *cvp2 cvl1* vascular phenotype seems to be PIN1-independent

To gain further insight into the mechanisms by which RPK2 modulates vein patterning we decided to monitor the transcript profiles of wild-type, *cvp2 cvl1* and *amiRPK2 cvp2 cvl1* embryos between the early torpedo and bent cotyledon stages. Both mutants showed hundreds of differentially expressed genes (DEGs) when compared to wild type (782 in the case of *cvp2 cvl1*, 423 in the case of *amiRPK2 cvp2 cvl1*). Remarkably, silencing *RPK2* expression resulted in a partial complementation at the transcriptomic level of the *cvp2 cvl1* defects (Fig. 6A-C), as evident for instance by the repression of the augmented transcript levels of *ALTERED PHLOEM (APL)* (Bonke et al. 2003) and *SISTER APL (SAPL)* (Ross-Elliott et al. 2017) of *cvp2 cvl1* embryos (Fig. 6D). Consistent with this result, the induced expression of At2g28810 (Furuta et al. 2014) or the callose biosynthetic enzyme *CALS7* (Vaten et al. 2011) *-*known targets of *APL-* got reverted to a wild-type situation in *amiRPK2 cvp2 cvl1* (Fig. 6D). Moreover, the expression of genes associated with mesophyll identity such as *CHLOROPHYLL A/B BINDING PROTEIN 3 (CAB3)* (Mitra et al. 1989) is not affected in *cvp2 cvl1* embryos (Fig. 6E), excluding an altered genetic program of mesophyll cells as responsible for *cvp2 cvl1* procambial cells’ misspecification. Surprisingly, our transcriptomic analysis did not reveal altered expression levels neither in auxin biosynthetic nor auxin related genes in *cvp2 cvl1* embryos (Fig. 6F), with the exception of the *AUXIN TRANSPORTER 1 (AUX1*) and *AUXIN TRANSPORTER-LIKE PROTEIN 1 (LAX1)* (Swarup and Bhosale 2019). Contrary to *cvp2 cvl1*, cotyledons of *aux1* and *lax1* null mutants exhibit a continuous vein network, even when both genes are simultaneously knocked-out (Fig. 6G, J-N). Introgression of *AUX1-GFP* transgene in *cvp2 cvl1* did not rescue the vascular phenotype of this mutant (Fig. 6G-I, M, N, Supporting Information Fig.1), inferring that *rpk2-*mediated rescue of *cvp2 cvl1* is not directly due to the modulation of auxin influx and its biosynthetic pathways. To exclude a potential regulation of PIN1-mediated auxin distribution by RPK2 as the restoring mechanism of *cvp2 cvl1* defects, we performed chemical treatments using a widely used auxin transport inhibitor, 1-naphtylphathalamic acid (NPA), in leaves. These organs form *de novo* during the post-embryonic growth of the plant. The establishment of the vascular strands in leaves is believed to follow a similar molecular regulation as cotyledons, even if the directionality of the secondary strands differs (Lavania et al. 2021). Under NPA treatment, the acropetal transport of auxin that directs the growth of the midvein is altered, resulting in the duplicated formation of midveins (Scarpella et al. 2006). Additionally, secondary vein formation is slightly affected whereas tertiary vein formation could appear suppressed (Scarpella et al. 2006). Similar to wild type and *cvp2 cvl1* plants, NPA-treated *rpk2* leaves appeared very affected in terms of vein complexity (Supporting Information Fig. S4A,F). Moreover, root growth inhibition of *rpk2* roots subjected to NPA treatments was very similar to the one observed in wild type and *cvp2 cvl1* plants (Supporting Information Fig.S4 L). These observations imply that RPK2 activity is not necessary to integrate auxin polar transport cues into the establishment of the vein pattern, despite the rescue of the *cvp2 cvl1* leaf vein pattern through silencing *RPK2* expression (Supporting Information Fig. S4G-I). Consistent with these results, live cell image analysis of PIN1-GFP localization in *rpk2* mutants did not reveal any defective PIN1 localization in this genetic background. Given the reduced offspring of *rpk2*, we decided to focus in the root stele to have a clear assessment of PIN1 polarity. Within the stele, PIN1 polarly accumulates at the basal cell membrane. The polar PIN1 distribution in *rpk2* roots (Supporting Information Fig. S4J-K’) together with the unaltered response of this mutant to NPA treatments suggests that RPK2 activity in modulating vein emergence is independent of PIN1-mediated auxin transport.

**Figure 6.**
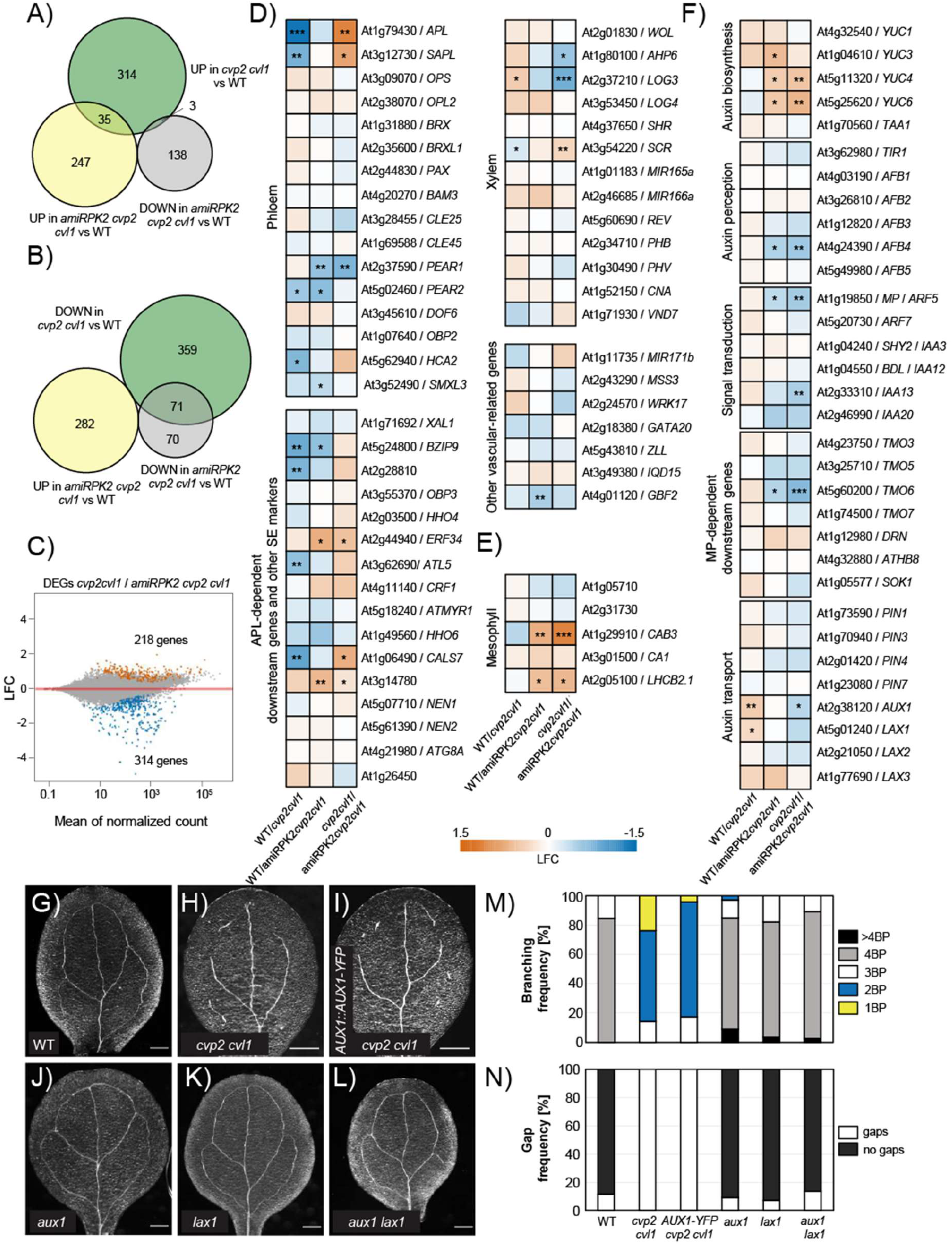
Differential gene expression analysis among WT, *cvp2 cvl1* and *amiRPK2 cvp2 cvl1*. **A-B)** Venn diagram showing the overlap of upregulated (A) and down-regulated (B) genes in *cvp2 cvl1* with the DEGs in *amiRPK2 cvp2 cvl1*. **C)** MA plot showing the log2 fold change (LFC) of each gene over the mean of normalized counts. **D-F)** Heatmaps show enrichment (LFC) of genes with known roles in vascular development (D), with known expression in mesophyll cells (E) and involved in auxin biosynthesis, signalling and transport (F). * represents p-value <0.05, ** represent p-value <0.01 and *** represent p-value <0.001. **G-L)** Bright-field images of 7-day-old cotyledons of the indicated genotypes. Scale bars represent 200μm. **M-N)** Quantification of gap and branching frequency of the vein network phenotypes observed in the cotyledons represented in G-L. n= 21-24 for each genotype. BP: branching points, counted as the initiation (even if not completed) of a new secondary vein.

### RPK2-mediated suppression of proximal branching is independent of CLE peptides

To further explore the underpinning molecular mechanisms of RPK2-mediated repression of secondary proximal veins, we analyzed the potential involvement of CLE peptides in controlling this process. *rpk2* loss-of-function mutant is resistant to the root growth-inhibition effect of a vast array of CLE peptides, including CLAVATA 3 (CLV3) and CLE45 (Gujas et al. 2020). In particular, RPK2 has been shown to perceive CLV3 peptide, negatively regulating root and shoot apical meristem cell proliferation (Racolta et al. 2018). Moreover, we have previously described that the *rpk2*-mediated restoration of a continuous root protophloem strand in *cvp2 cvl1* involves CLE45 (Gujas et al. 2020). To investigate the potential role of both peptides in orchestrating cotyledon vascular patterning during embryogenesis, we cleared cotyledons of *clv3* and *cle45* mutant seedlings (Yamaguchi et al. 2017). Neither *cle45* nor *clv3* cotyledons showed any perturbation in their vein pattern or vein network complexity (Supporting Information Fig. S5A-D, G), suggesting that they are either not involved in regulating cotyledon vein networks or that there is a high redundancy within this family of peptides. An overexpression of *CLE45* results in lethality (Depuydt et al. 2013). Thus, we aimed at increasing *CLE45* and *CLV3* expression specifically during embryogenesis. To this aim we generated transgenic lines expressing *CLE45* and *CLV3* under the *ABSCISIC ACID INSENSITIVE 3 (ABI3)* promoter. *ABI3* is broadly expressed in the embryo, beginning at the globular stage until embryo maturation and ceases to be expressed after germination (To et al. 2006). Consistent with the previous results (Supporting Information Fig. S5), analysis of 8-day old *ABI3∷CLE45* seedlings did not reveal any vascular defects in the cotyledons, implying that RPK2-mediated regulation of vascular patterning is independent of CLE45 (Supporting Information Fig. S5E, G). In contrast, a discontinuous vascular network in 25% of *ABI3∷CLV3* seedlings was detected, even if this frequency is 5% in wild type plants (Supporting Information Fig. S5F, G). This feature was associated with reduced proximal branching and, in turn, vascular complexity (Supporting Information Fig. S5F, G). Our results indicated that despite this peptide having the ability to supress proximal vein branching, its restricted expression at the shoot apical meristem (Brand et al. 2002) most likely excludes it as the ligand responsible for RPK2-mediated control of proximal branching.

### OCTOPUS promotes proximal branching in embryonic cotyledons

Another factor involved in the regulation of procambial cell division in embryonic cotyledons is the plasma membrane-associated protein OCTOPUS (OPS) (Truernit et al. 2012). In *ops* embryos, a reduced number of cell files in distal secondary veins can be detected as the procambial cells fail to periclinally divide (Roschzttardtz et al. 2014). Live-cell imaging analysis of OPS distribution revealed a broader expression domain than that of the future vein path in intermediate torpedo stage embryos (Fig. 7I-I’) and a non-polar cellular distribution in cells not within the vein path (Fig. 7J, J’). In contrast, OPS appears polarly distributed in the apical cell membrane of procambial cells in intermediate torpedo stage embryos and in mature embryos once the vein network has achieved its final complexity (Fig. 7K-L’,M). OPS distribution can also be detected in the non-procambial cells adjacent to the branching point and close to the cotyledon margin of intermediate torpedo embryos, near the *RPK2* expression domain (Fig. 7I-J). Interestingly, an increase in OPS protein dosage by introducing the hyperactive GFP-tagged OPS protein (Supporting Information Fig. S1) (Breda et al. 2017) increased the number of proximal branching points in wild type plants (Fig. 7A, C, F). Likewise, an increase in branching points was observed in *cvp2 cvl1* as well as an overall more continuous cotyledon vein network, even though vein pattern discontinuities still persisted (Fig. 7A-H). These results confirmed that CVP2 and CVL1 are necessary to establish the proximal branching of secondary veins, a process enhanced by the positive vascular regulator OPS and counteracted by RPK2.

**Figure 7.**
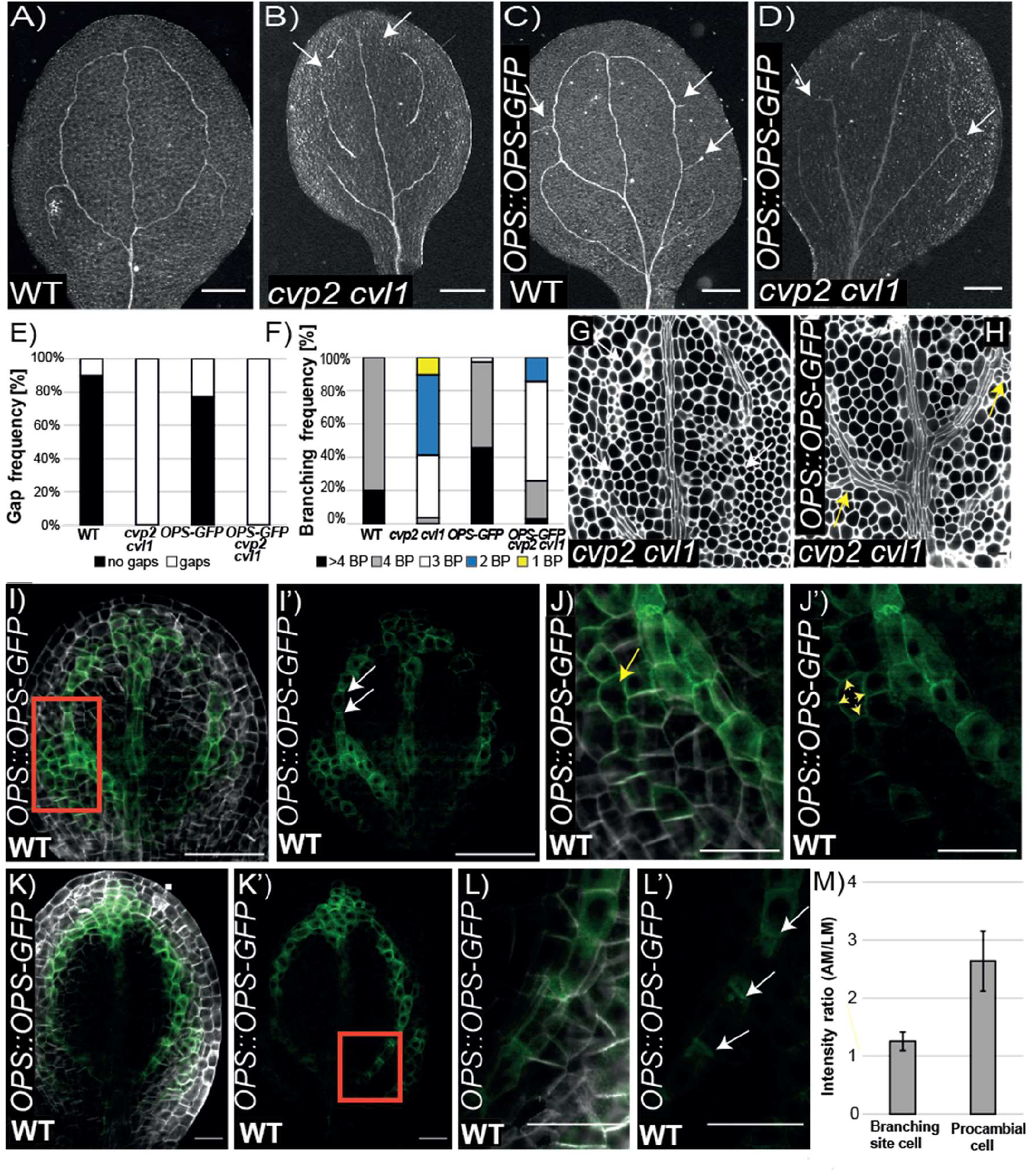
OCTOPUS promotes proximal branching in WT and *cvp2 cvl1* embryonic cotyledons. **A-D)** Representative images of cleared 8-day-old cotyledons of the indicated genotypes imaged with a stereomicroscope in bright field on a black background. White arrows mark vein gaps and yellow arrows indicate additional proximal branching sites. Scale bars represent 200μm. **E-F)** Quantification of gap and branching frequency of the vein network phenotypes observed in the cotyledons represented in A-J. n= 29-35 for each genotype. BP: branching points, counted as the initiation (even if not completed) of a new secondary vein. **G-H)** mPS-PI stained embryos visualized by confocal microscopy of the indicated genotypes. Scale bars represent 20 μm**. I-L’)** Visualization of OPS distribution in early (I-J’) and late (K-L’) torpedo stage embryos stained with Renaissance. J and L represents a magnification of the branching region represented in I and K. In I’, J’, K’ and L’ only the GFP signal is shown. Scale bars represent 50 μm in, I, I’, K, K’ and 20 μm in J, J’, L, L’. **M)** Quantification of OPS polarity in cells from early and late torpedo stages as means of the ratio of GFP signal between the apical and lateral membrane.

## Discussion

### CVP2 and CVL1 regulate the specification of provascular cells and proximal branching

Within a multicellular organ, the self-establishment of tissue patterns such as vascular tissues requires the coordinated activity of oriented cell divisions and cell fate commitment (Lavania et al. 2021). Our data reveal that *CVP2* and *CVL1* contribute to the regulation of vein patterning by modulating vascular specification and the branching points of proximal secondary veins (Fig. 2). On the one hand, *cvp2 cvl1* cotyledons exhibit a discontinuous vein network because of GM cell misspecification into mesophyll instead of pro-vascular cells (Fig. 2). A defective vascular identity perfectly matches with the lack of auxin activity in these cells, as revealed by auxin biosensors (Fig. 4L, M). Yet, the lack of this hormone in these cells does not correlate with a significant different auxin transcriptional response in *cvp2 cvl1* embryos at an earlier developmental stage, the torpedo stage (Fig. 6). Moreover, our analysis of vascular identity domains in *cvp2 cvl1* globular and heart stage embryos did not reveal defects in the establishment of vascular *vs* ground tissue domains over embryogenesis (Supporting Information Fig. S2). Thus, an alternative explanation consistent with *cvp2 cvl1* phenotypes is that a premature differentiation of mesophyll cells terminates vein propagation, a phenomenon that has been described in leaves (Scarpella et al. 2004). However, further experiments are required to elucidate whether *CVP2* and *CVL1* participate in repressing mesophyll differentiation, even if the transcripts associated to mesophyll identity did not appear significantly altered in *cvp2 cvl1* embryos as evident by our transcriptomic profiling (Fig. 6E). Alternatively, the deficient phloem-mediated sugar transport in *cvp2 cvl1* cotyledons (Rodriguez-Villalon et al. 2015; Carland and Nelson 2009) may inhibit the regulation of GM specification into procambial cells. Further studies are required to assess whether sucrose intervenes in the conversion of GM into either mesophyll or procambial cells and whether a perturbed sucrose distribution may translate in the appearance of vascular islands in *cvp2 cvl1*. While these disconnected vascular islands can be frequently observed in the *cvp2 cvl1* cotyledon vein network, the midvein always appears intact (Fig. 2). In cotyledons, the midvein originates directly from a pool of cells located in the vicinity of the shoot apical meristem, which start to elongate and divide parallel to the proximo-distal axis of cotyledon growth at the torpedo stage (Nelson and Dengler 1997). This process is mostly regulated by auxin signalling factors such as *ARF6/CULLIN1* or *ARF5/MONOPTEROS* (Scarpella et al. 2004; Scarpella et al. 2006), implying that the activity of these factors remains unaffected by the loss of *CVP2* and *CVL1* during embryogenesis until early torpedo stage. Several vascular-associated genes appeared miss-expressed in *cvp2 cvl1* embryos, even if *RPK2* silencing only reverted *APL* (and APL- downstream target genes) as well as *SAPL* expression (Fig. 6). On the other hand, the reduced vein network complexity described in *cvp2 cvl1* cotyledons reflects the inability of proximal secondary veins to initiate branching. Together with the aberrant divisions detected in the suspensor and hypophysis (Supporting Information Fig. S3), these observations infer that the activity of both CVP2 and CVL1 may be required to a certain extent, to control the orientation of cell divisions. Interestingly, closer examination at the branching point in wild type plants revealed a periclinal division of a distal vein cell next to another cell harbouring either a vascular marker or *DR5* (Supplemental Information Fig. S6). While it appears possible that a periclinal division precedes the formation of a vascular cells from which the incipient proximal secondary vein will extend, we cannot exclude at this stage that a plate meristematic cell adjacent to the vascular cells is recruited at the branching point to give rise to the new vein. Another interesting aspect of this process is whether auxin is involved in the coordination of proximal vein branching. Genetic blockage of polar auxin transport by depleting the activity of PIN transporter results in an elevated number of distal veins with additional distal branching points (Fig. 4). While these results indicated that polar auxin cues contribute to repress distal branching, they cannot explain the reduced cotyledon vein network complexity observed in *cvp2 cvl1* nor its intermittent cotyledon vein pattern. Our observations show that PIN1 polarity appears unaltered in *cvp2 cvl1* veins (Fig. 3). Additionally, genetic increase of PIN1 dosage does not enhance the branching defects observed in *cvp2 cvl1* (Fig. 3), indicating that at least another mechanism independent of PIN1 must be responsible for this phenotype. Previous studies have shown that regardless of the absence of auxin transport, the remnant auxin signalling is sufficient to guide the recruitment of new procambial cells into vascular cell files (Verna et al. 2019). However, auxin response in *cvp2 cvl1* cotyledons by means of *DR5* distribution appears discontinuous in the vein path. Hence, it appears possible that an auxin-independent mechanism preceding *PIN1* expression is required to modulate vascular cell identity acquisition and in turn, the future vein path.

### *RPK2* constrains proximal secondary vein branching

In globular embryos, *RPK2* is expressed in the outermost layer that gives rise to the ground tissues, an expression pattern that is maintained during the following developmental stages of embryogenesis (Fig. 5) (Nodine et al. 2007). Previous studies have suggested that *RPK2* is required to exclude vascular identity from the protodermis of globular embryos (Nodine et al. 2007). Here, we provide evidence that RPK2 activity is not only necessary to exclude vascular identity in the future epidermis but also necessary to modulate the branching of secondary veins (Fig. 5). The activity of RPK2 in controlling the development of cortical and endodermal cells have been mostly explained by the perception of CLE 17 (Racolta et al. 2018). Within the root stele, CLE45 sensing by RPK2 confers developmental plasticity to companion and protophloem cells to safeguard phloem functionality by re-establishing a correct phloem pattern in case the original one fails to form (Gujas et al. 2020). Yet, our analysis implies that the suppression of cotyledon proximal branching by RPK2 is independent of vascular-specific CLE peptides (Supporting Information Fig. S5), inferring a differential mechanism between vascular formation in the shoot and the root. Our results indicate that the negative RPK2-mediated control of vein patterning is independent of auxin. Yet, plant cells need to integrate PAT cues and RPK2 signalling to coordinate the establishment and maintenance of vascular tissues. Considering that *rpk2* sensitivity to the blockage of PAT by NPA is not perturbed (Supporting Information Fig. S4), additional factors other than RPK2 may act as a hub to integrate polar auxin transport cues with RPK2-mediated signalling. *RPK2* expression ends at the cell file delineating the border between the cotyledon margin and the vein cells, in a region where the positive vascular regulator OPS is localized before the establishment of a closed vein network (Fig. 5L, L’, 7I-K). While future studies will elucidate whether the mutually exclusive presence of OPS and RPK2 is required to trigger proximal vein branching, it appears possible that OPS contributes to integrate PAT cues with RPK2 signalling by means of its interaction with VASCULATURE COMPLEXITY AND CONNECTIVITY (VCC) (Roschzttardtz et al. 2014). The latter contributes to spatio-temporally modulate PAT during cotyledon vein formation by delivering PIN1 to the vacuole for protein degradation (Yanagisawa et al. 2021). Together, our results revealed a molecular genetic framework by which plants at the late stages of embryogenesis modulate the tissue complexity of their vascular networks. Although the function of *RPK2* is conserved in cylindrical and foliar organs, its regulation appears to be tissue specific, comprising unique molecular mechanisms to optimize the functionality of vascular tissues to the constantly changing shape of the organs to which they belong.

## Acknowledgements

We greatly thank Dr. Dolf Weijers, Margo Smit and Thijs de Zeeuw for technical assistance with imaging embryos. Dr. Jiri Friml, Dr. Takashi Ishida, Dr. Olivier Hamant, Dr. Enrico Scarpella, Dr.Miguel A. Perez Amador, Dr. Ari Pekka Mahonen and Dr. Christian Fankhauser for kindly providing seeds and Functional Genomic Center of Zurich and ScopeM for their services. E.K. was funded by the Stravros Niarchos-ETH Foundation. This work was funded by the Swiss National Foundation (SNF_31003A_160201 to A.R.-V.).

## Author contributions

AR-V, EK and NB-T designed the research experiments. EK, NB-T, AS performed experiments and data analysis. AS and BG generated genetic material. AR-V, EK and NB-T wrote the manuscript.

**Supporting Information Fig Supplemental 1.** Transcriptional profiling of the transgenic lines used in the current study.

**Supporting Information Fig. Supplemental 2.** Analysis of *CVP2* and *CVL1* expression and vascular identity domains in *cvp2 cvl1* embryos.

**Supporting Information Fig. Supplemental 3.** *cvp2 cvl1* embryos exhibit aberrant divisions.

**Supporting Information Fig. Supplemental 4.** Sensitivity to PIN1-mediated auxin transport is not disturbed in *rpk2*.

**Supporting Information Fig. Supplemental 5.** RPK2 modulation of vascular branching is independent of vascular-specific CLE peptides.

**Supporting Information Fig. Supplemental 6.** A periclinal division occurs at the branching point.

---

**Fig. S1.**
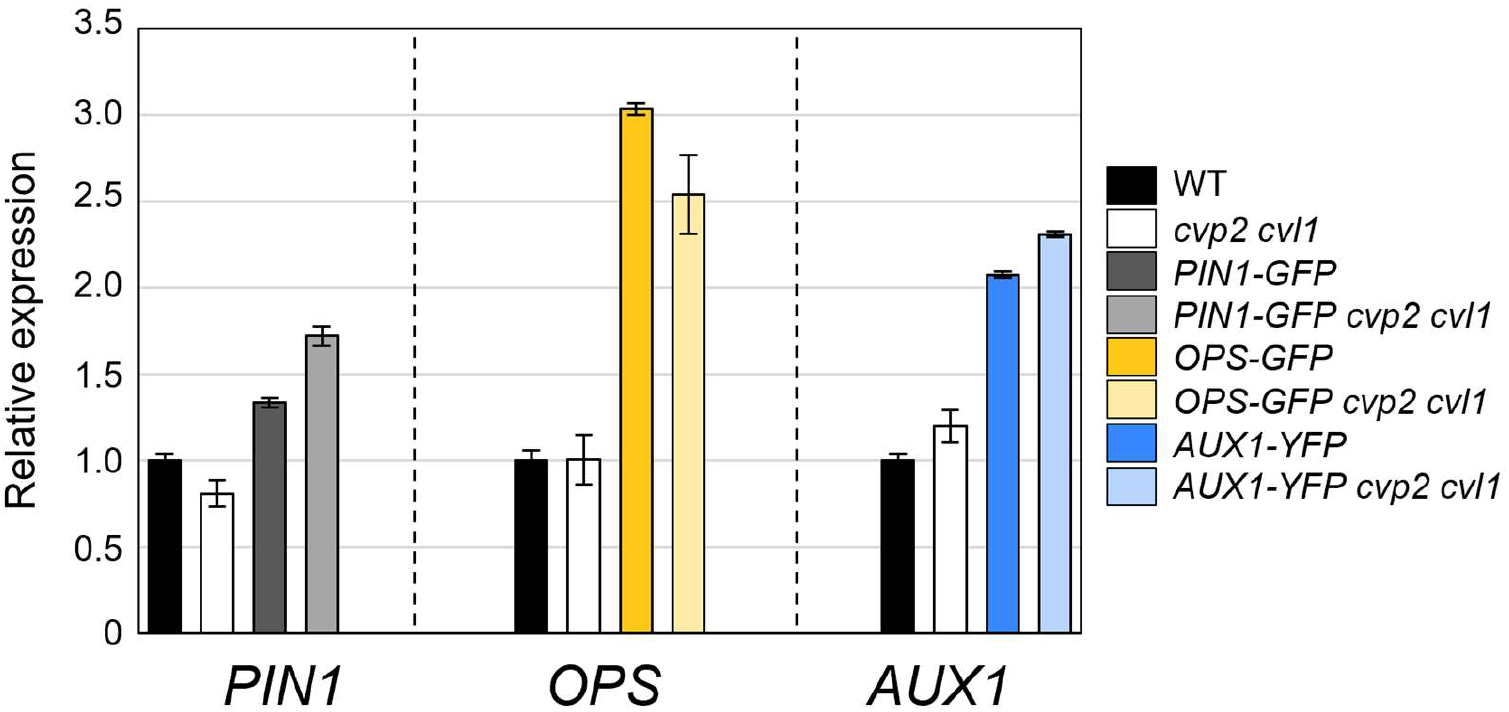
Transcriptional profiling of the transgenic lines used in the current study. Levels of *PIN1*, *OPS* and *AUX1* normalized expression in 7-day-old seedlings of the indicated genotypes in comparison to wild type. Values represent the mean of 3 technical replicates and error bars represent the standard deviation of these replicates.

**Fig. S2.**
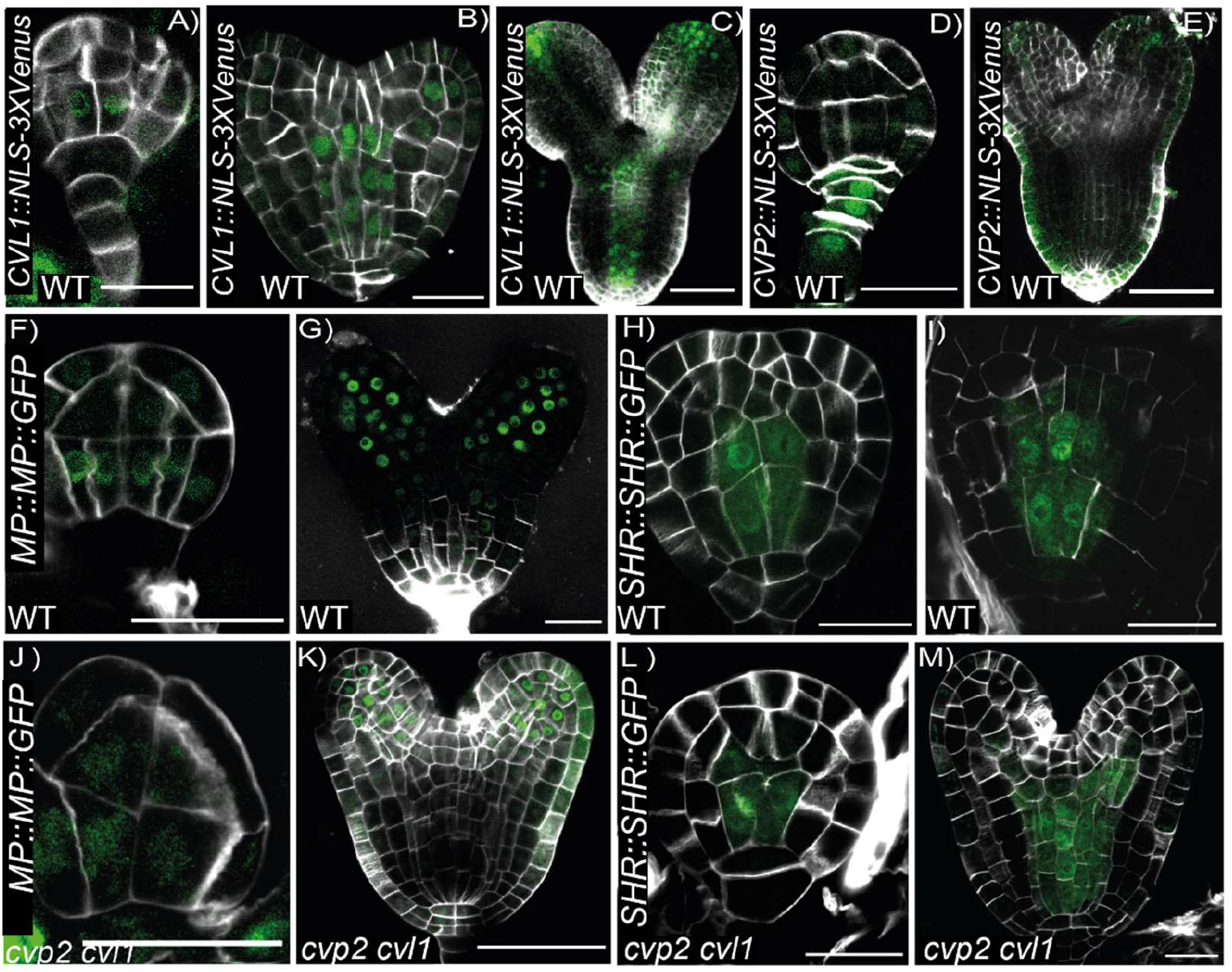
Analysis of *CVP2* and *CVL1* expression and vascular identity domains in *cvp2 cvl1* embryos. **A-E)** Expression pattern of the indicated genes in early globular (A, D), heart (B, E) and torpedo (C) developmental stages. Renaissance SR2200 staining highlights patterning divisions by labelling plant cell walls. Scale bars represent 50μm in A, C, D, E and 20μm in B. **(F-I)** Expression pattern of the indicated vascular genes in globular (F, H, J, L) and heart stage (G, I, K, M) embryos of WT and *cvp2 cvl1*. Scale bars represent 20μm in (G, I, K, M) and 50μm in (F, H, J, L).

**Fig. S3.**
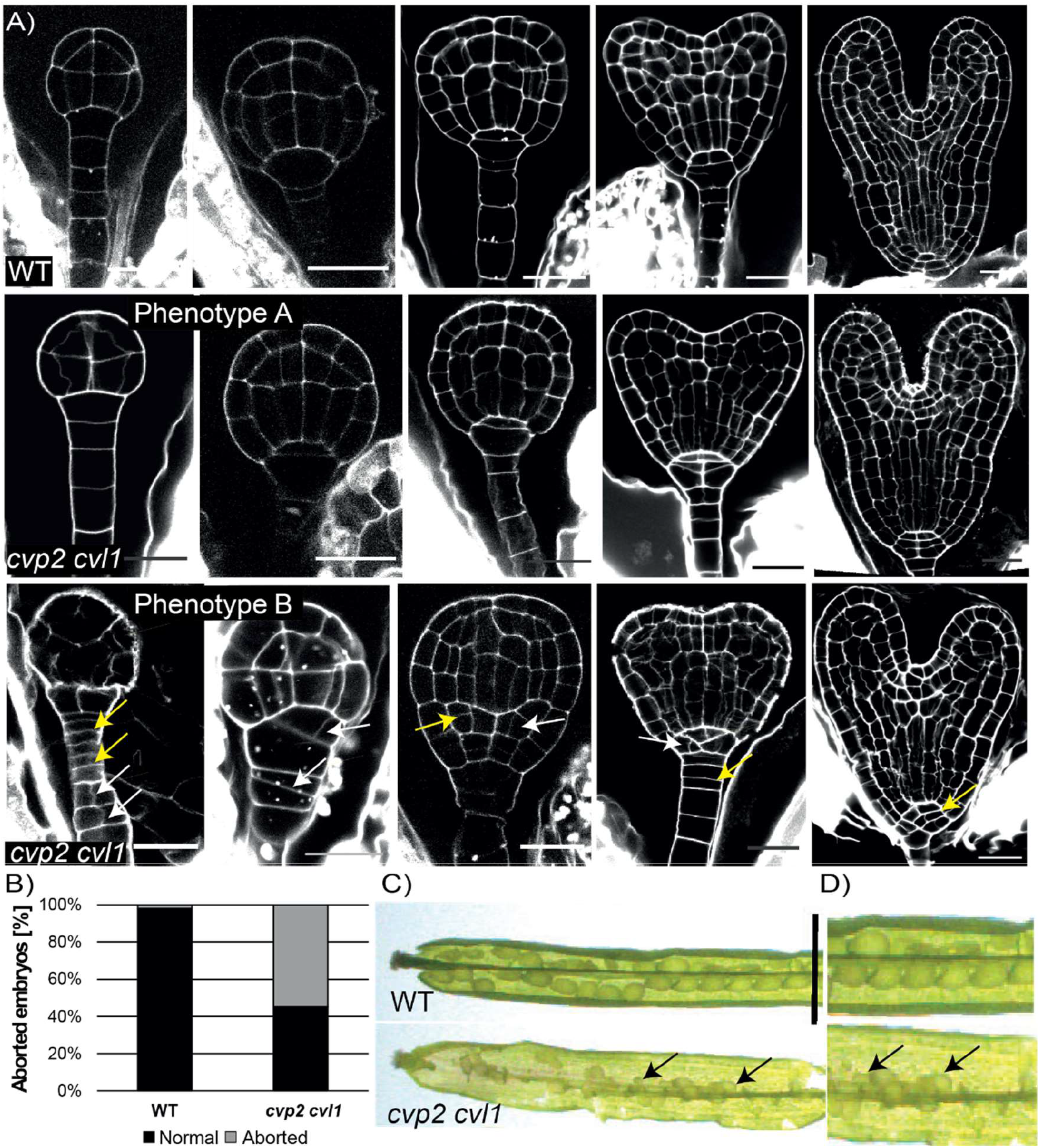
*cvp2 cvl1* embryos exhibit aberrant divisions. **A)** Embryo morphology of WT compared to *cvp2 cvl1* embryos using mSP-PI on ovules taken from green siliques and imaged using confocal microscopy. *cvp2 cvl1* shows aberrant divisions at all stages of embryogenesis but most abundantly during the globular stage. Yellow arrows point to aberrant divisions, and scale bars are 20μm or 50μm. **B)** Graphical representation of the percent of aborted embryos occurring in *cvp2 cvl1* vs. WT siliques numbered 2 and 3 counted from the apical meristem (containing globular stage embryos). **C)** Image of a dissected WT silique as compared to *cvp2 cvl1* under bright field using a stereomicroscope. Scale bar represents 1inch.

**Fig. S4.**
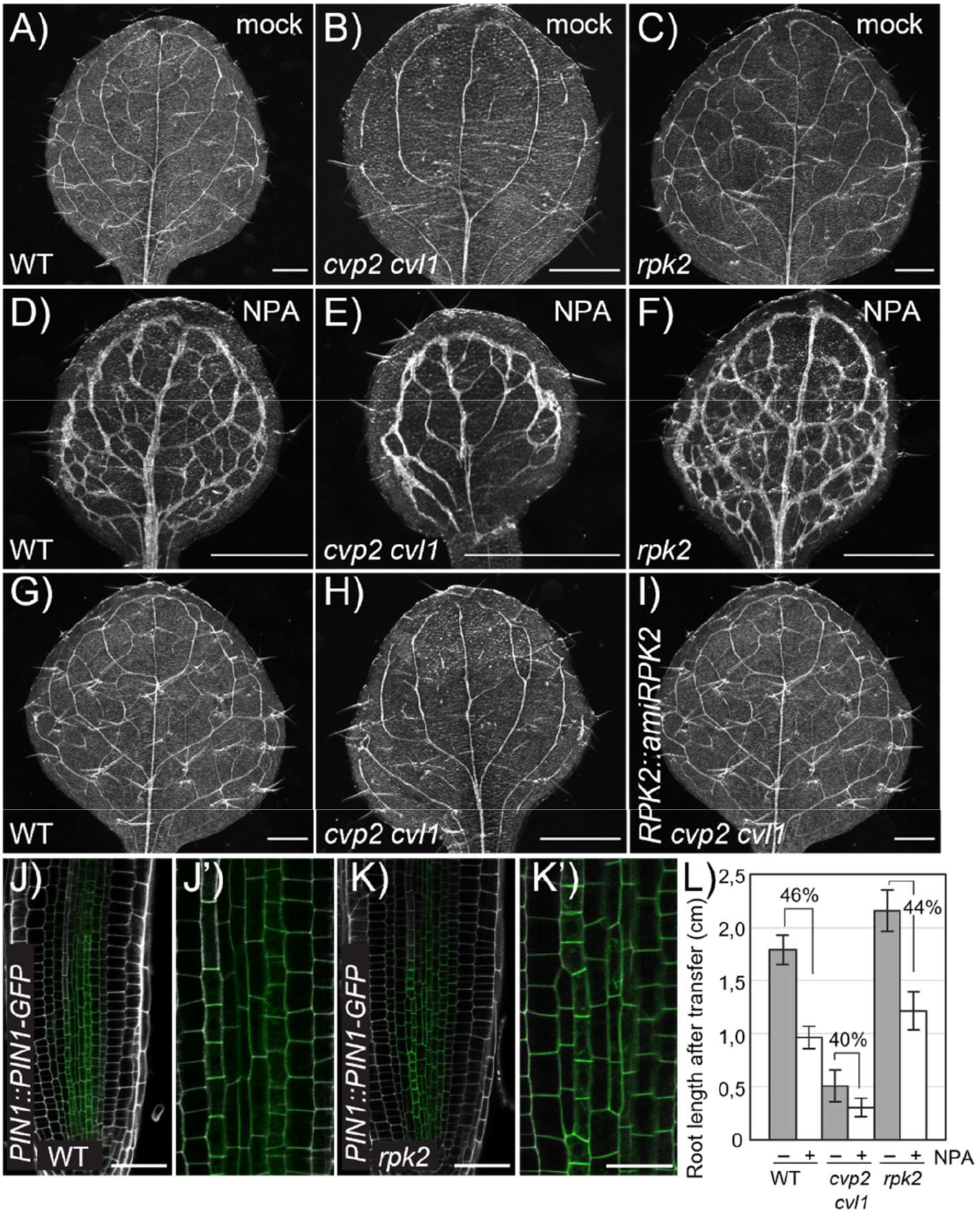
Sensitivity to PIN1-mediated auxin transport is not disturbed in *rpk2*. **A-F)** Analysis of cleared leaves of seedlings grown for 4 days (until clear emergence of cotyledons could be detected) transferred to a media supplemented with 10 μM NPA or mock conditions for 5 days. n= 19-35 for each genotype. Scale bars: 500 μm. **G-I)** Cleared leaves of the indicated genotypes imaged with a stereomicroscope in bright field on a black background. n= 14-21 for each genotype. **J-K’)** Confocal microscopy analysis of *PIN1∷PIN1-GFP* distribution in cells of the root stele of WT and *rpk2-2* seedlings. J’ and K’ represent magnification of the region shown in J and K. Scale bars represent 50μm in J,K and 20μm in J’ and K’. **L)** Root length of seedlings grown as described in A-F) were measured. Note that root length was measured after the treatment with NPA or mock. The root length of seedlings treated with mock were set to 100% and the % shown in the graph represents the % of inhibition of root length by NPA. n=39-56.

**Fig. S5.**
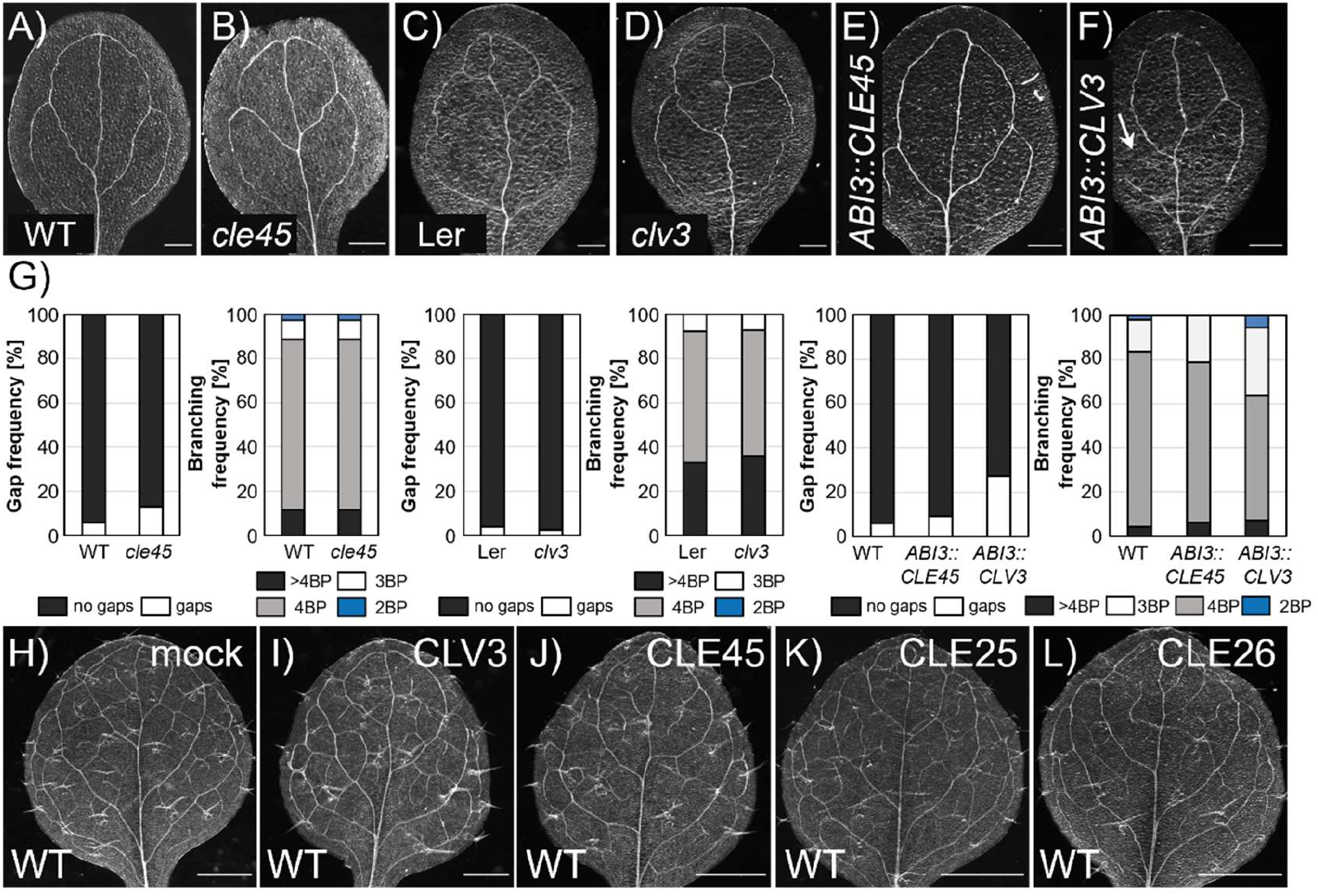
RPK2 modulation of vascular branching is independent of vascular-specific CLE peptides. **A-F)** Analysis of the continuity and complexity of cotyledon vein network in 8-day-old seedlings of the indicated genetic backgrounds. Arrow in F indicates defects in proximal branching in *pABI3∷CVL3*. Scale bars represent 200μm. **G)** Quantification of the gap and branching frequency in the cotyledons analyzed in A-F. n= 17-55 for each genotype. **H-L)** Analysis of the vein pattern in cleared leaves of 9-day-old seedlings transferred to a medium supplemented with the indicated CLE peptides once the emergence of the cotyledons could be detected. n= 16-46 for each genotype. Scale bars represent 500μm.

**Fig. S6.**
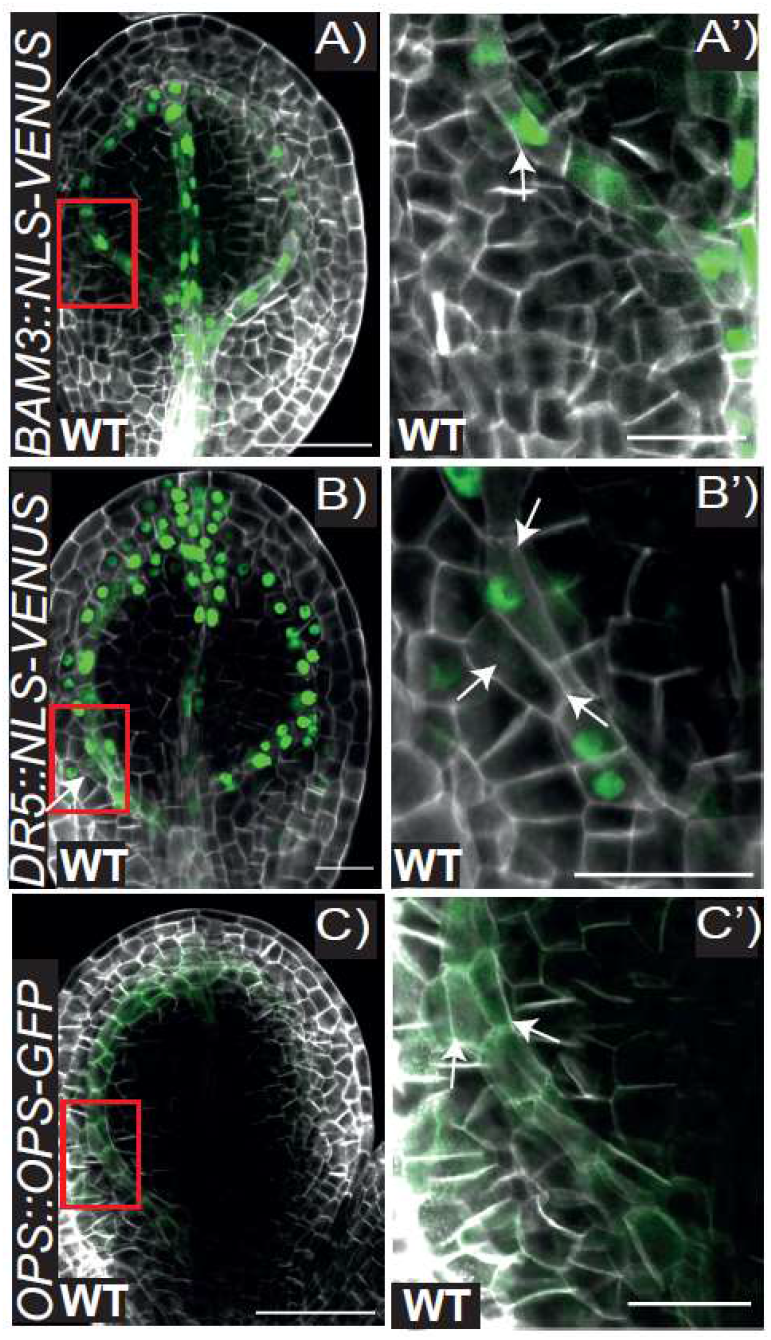
A periclinal division occurs at the branching point. Representative pictures of early torpedo stage embryos harbouring *BAM3∷NLS-3xVENUS* (A), *DR5∷NLS-VENUS* (B) *and OPS∷OPS-GFP* (C). Embryonic cotyledons were stained with Renaissance stain SR2200 and visualized by confocal microscopy. Magnification of the branching region squared in A), B) and C) is shown in A’), B’) and C’), respectively. White arrows indicate nuclei in the cells having undergone a periclinal cell division at the branching point. Scale bars represent 50μm in A,C and 20μm A’-C’,B.

## References

Agusti J, Blazquez MA (2020) Plant vascular development: mechanisms and environmental regulation. Cell Mol Life Sci. doi:10.1007/s00018-020-03496-w

Bonke M, Thitamadee S, Mahonen AP, Hauser MT, Helariutta Y (2003) APL regulates vascular tissue identity in Arabidopsis. Nature 426 (6963):181–186. doi:10.1038/nature02100

Brand U, Grunewald M, Hobe M, Simon R (2002) Regulation of CLV3 expression by two homeobox genes in Arabidopsis. Plant Physiol 129 (2):565–575. doi:10.1104/pp.001867

Breda AS, Hazak O, Hardtke CS (2017) Phosphosite charge rather than shootward localization determines OCTOPUS activity in root protophloem. Proc Natl Acad Sci U S A 114 (28):E5721–E5730. doi:10.1073/pnas.1703258114

Carland F, Nelson T (2009) CVP2- and CVL1-mediated phosphoinositide signaling as a regulator of the ARF GAP SFC/VAN3 in establishment of foliar vein patterns. Plant J 59 (6):895–907. doi:10.1111/j.1365-313X.2009.03920.x

Carland FM, Nelson T (2004) Cotyledon vascular pattern2-mediated inositol (1,4,5) triphosphate signal transduction is essential for closed venation patterns of Arabidopsis foliar organs. Plant Cell 16 (5):1263–1275. doi:10.1105/tpc.021030

Czechowski T, Stitt M, Altmann T, Udvardi MK, Scheible WR (2005) Genome-wide identification and testing of superior reference genes for transcript normalization in Arabidopsis. Plant Physiology 139 (1):5–17. doi:10.1104/pp.105.063743

De Rybel B, Mahonen AP, Helariutta Y, Weijers D (2016) Plant vascular development: from early specification to differentiation. Nat Rev Mol Cell Bio 17 (1). doi:10.1038/nrm.2015.6

Depuydt S, Rodriguez-Villalon A, Santuari L, Wyser-Rmili C, Ragni L, Hardtke CS (2013) Suppression of Arabidopsis protophloem differentiation and root meristem growth by CLE45 requires the receptor-like kinase BAM3. Proc Natl Acad Sci U S A 110 (17):7074–7079. doi:10.1073/pnas.1222314110

Dindas J, Scherzer S, Roelfsema MRG, von Meyer K, Muller HM, Al-Rasheid KAS, Palme K, Dietrich P, Becker D, Bennett MJ, Hedrich R (2018) AUX1-mediated root hair auxin influx governs SCF(TIR1/AFB)-type Ca(2+) signaling. Nat Commun 9 (1):1174. doi:10.1038/s41467-018-03582-5

Friml J, Vieten A, Sauer M, Weijers D, Schwarz H, Hamann T, Offringa R, Jurgens G (2003) Efflux-dependent auxin gradients establish the apical-basal axis of Arabidopsis. Nature 426 (6963):147–153. doi:10.1038/nature02085

Furuta KM, Yadav SR, Lehesranta S, Belevich I, Miyashima S, Heo JO, Vaten A, Lindgren O, De Rybel B, Van Isterdael G, Somervuo P, Lichtenberger R, Rocha R, Thitamadee S, Tahtiharju S, Auvinen P, Beeckman T, Jokitalo E, Helariutta Y (2014) Plant development. Arabidopsis NAC45/86 direct sieve element morphogenesis culminating in enucleation. Science 345 (6199):933–937. doi:10.1126/science.1253736

Gujas B, Cruz TMD, Kastanaki E, Vermeer JEM, Munnik T, Rodriguez-Villalon A (2017) Perturbing phosphoinositide homeostasis oppositely affects vascular differentiation in Arabidopsis thaliana roots. Development 144 (19):3578–3589. doi:10.1242/dev.155788

Gujas B, Kastanaki E, Sturchler A, Cruz TMD, Ruiz-Sola MA, Dreos R, Eicke S, Truernit E, Rodriguez-Villalon A (2020) A Reservoir of Pluripotent Phloem Cells Safeguards the Linear Developmental Trajectory of Protophloem Sieve Elements. Curr Biol 30 (5):755–766 e754. doi:10.1016/j.cub.2019.12.043

Gujas B, Rodriguez-Villalon A (2016) Plant Phosphoglycerolipids: The Gatekeepers of Vascular Cell Differentiation. Front Plant Sci 7:103. doi:10.3389/fpls.2016.00103

Heisler MG, Ohno C, Das P, Sieber P, Reddy GV, Long JA, Meyerowitz EM (2005) Patterns of auxin transport and gene expression during primordium development revealed by live imaging of the Arabidopsis inflorescence meristem. Curr Biol 15 (21):1899–1911. doi:10.1016/j.cub.2005.09.052

Lavania D, Linh NM, Scarpella E (2021) Of Cells, Strands, and Networks: Auxin and the Patterned Formation of the Vascular System. Cold Spring Harb Perspect Biol. doi:10.1101/cshperspect.a039958

Lucas WJ, Groover A, Lichtenberger R, Furuta K, Yadav SR, Helariutta Y, He XQ, Fukuda H, Kang J, Brady SM, Patrick JW, Sperry J, Yoshida A, Lopez-Millan AF, Grusak MA, Kachroo P (2013) The plant vascular system: evolution, development and functions. J Integr Plant Biol 55 (4):294–388. doi:10.1111/jipb.12041

Marhava P, Aliaga Fandino AC, Koh SWH, Jelinkova A, Kolb M, Janacek DP, Breda AS, Cattaneo P, Hammes UZ, Petrasek J, Hardtke CS (2020) Plasma Membrane Domain Patterning and Self-Reinforcing Polarity in Arabidopsis. Dev Cell 52 (2):223–235 e225. doi:10.1016/j.devcel.2019.11.015

Mazur E, Kulik I, Hajny J, Friml J (2020) Auxin canalization and vascular tissue formation by TIR1/AFB-mediated auxin signaling in Arabidopsis. New Phytol 226 (5):1375–1383. doi:10.1111/nph.16446

Mitra A, Choi HK, An G (1989) Structural and functional analyses of Arabidopsis thaliana chlorophyll a/b-binding protein (cab) promoters. Plant Mol Biol 12 (2):169–179. doi:10.1007/BF00020502

Naramoto S, Sawa S, Koizumi K, Uemura T, Ueda T, Friml J, Nakano A, Fukuda H (2009) Phosphoinositide-dependent regulation of VAN3 ARF-GAP localization and activity essential for vascular tissue continuity in plants. Development 136 (9):1529–1538. doi:10.1242/dev.030098

Nelson T, Dengler N (1997) Leaf Vascular Pattern Formation. Plant Cell 9 (7):1121–1135. doi:10.1105/tpc.9.7.1121

Nodine MD, Yadegari R, Tax FE (2007) RPK1 and TOAD2 are two receptor-like kinases redundantly required for arabidopsis embryonic pattern formation. Dev Cell 12 (6):943–956. doi:10.1016/j.devcel.2007.04.003

Palovaara J, Saiga S, Wendrich JR, Hofland NV, van Schayck JP, Hater F, Mutte S, Sjollema J, Boekschoten M, Hooiveld GJ, Weijers D (2017) Transcriptome dynamics revealed by a gene expression atlas of the early Arabidopsis embryo. Nature Plants 3 (11):894–904. doi:10.1038/s41477-017-0035-3

Przemeck GK, Mattsson J, Hardtke CS, Sung ZR, Berleth T (1996) Studies on the role of the Arabidopsis gene MONOPTEROS in vascular development and plant cell axialization. Planta 200 (2):229–237. doi:10.1007/BF00208313

Racolta A, Nodine MD, Davies K, Lee C, Rowe S, Velazco Y, Wellington R, Tax FE (2018) A Common Pathway of Root Growth Control and Response to CLE Peptides Through Two Receptor Kinases in Arabidopsis. Genetics 208 (2):687–704. doi:10.1534/genetics.117.300148

Rodriguez-Villalon A, Gujas B, Kang YH, Breda AS, Cattaneo P, Depuydt S, Hardtke CS (2014) Molecular genetic framework for protophloem formation. Proc Natl Acad Sci U S A 111 (31):11551–11556. doi:10.1073/pnas.1407337111

Rodriguez-Villalon A, Gujas B, van Wijk R, Munnik T, Hardtke CS (2015) Primary root protophloem differentiation requires balanced phosphatidylinositol-4,5-biphosphate levels and systemically affects root branching. Development 142 (8):1437–1446. doi:10.1242/dev.118364

Roschzttardtz H, Paez-Valencia J, Dittakavi T, Jali S, Reyes FC, Baisa G, Anne P, Gissot L, Palauqui JC, Masson PH, Bednarek SY, Otegui MS (2014) The VASCULATURE COMPLEXITY AND CONNECTIVITY gene encodes a plant-specific protein required for embryo provasculature development. Plant Physiol 166 (2):889–902. doi:10.1104/pp.114.246314

Ross-Elliott TJ, Jensen KH, Haaning KS, Wager BM, Knoblauch J, Howell AH, Mullendore DL, Monteith AG, Paultre D, Yan D, Otero S, Bourdon M, Sager R, Lee JY, Helariutta Y, Knoblauch M, Oparka KJ (2017) Phloem unloading in Arabidopsis roots is convective and regulated by the phloem-pole pericycle. Elife 6. doi:10.7554/eLife.24125

Scarpella E (2017) The logic of plant vascular patterning. Polarity, continuity and plasticity in the formation of the veins and of their networks. Curr Opin Genet Dev 45:34–43. doi:10.1016/j.gde.2017.02.009

Scarpella E, Francis P, Berleth T (2004) Stage-specific markers define early steps of procambium development in Arabidopsis leaves and correlate termination of vein formation with mesophyll differentiation. Development 131 (14):3445–3455. doi:10.1242/dev.01182

Scarpella E, Marcos D, Friml J, Berleth T (2006) Control of leaf vascular patterning by polar auxin transport. Genes Dev 20 (8):1015–1027. doi:10.1101/gad.1402406

Smit ME, Llavata-Peris CI, Roosjen M, van Beijnum H, Novikova D, Levitsky V, Sevilem I, Roszak P, Slane D, Jurgens G, Mironova V, Brady SM, Weijers D (2020) Specification and regulation of vascular tissue identity in the Arabidopsis embryo. Development 147 (8). doi:ARTN dev186130 10.1242/dev.186130

Swarup R, Bhosale R (2019) Developmental Roles of AUX1/LAX Auxin Influx Carriers in Plants. Front Plant Sci 10:1306. doi:10.3389/fpls.2019.01306

ten Hove CA, Lu KJ, Weijers D (2015) Building a plant: cell fate specification in the early Arabidopsis embryo. Development 142 (3):420–430. doi:10.1242/dev.111500

To A, Valon C, Savino G, Guilleminot J, Devic M, Giraudat J, Parcy F (2006) A network of local and redundant gene regulation governs Arabidopsis seed maturation. Plant Cell 18 (7):1642–1651. doi:10.1105/tpc.105.039925

Truernit E, Bauby H, Belcram K, Barthelemy J, Palauqui JC (2012) OCTOPUS, a polarly localised membrane-associated protein, regulates phloem differentiation entry in Arabidopsis thaliana. Development 139 (7):1306–1315. doi:10.1242/dev.072629

Truernit E, Bauby H, Dubreucq B, Grandjean O, Runions J, Barthelemy J, Palauqui JC (2008) High-resolution whole-mount imaging of three-dimensional tissue organization and gene expression enables the study of Phloem development and structure in Arabidopsis. Plant Cell 20 (6):1494–1503. doi:10.1105/tpc.107.056069

Tsukaya H (2021) The leaf meristem enigma: The relationship between the plate meristem and the marginal meristem. Plant Cell. doi:10.1093/plcell/koab190

Vaten A, Dettmer J, Wu S, Stierhof YD, Miyashima S, Yadav SR, Roberts CJ, Campilho A, Bulone V, Lichtenberger R, Lehesranta S, Mahonen AP, Kim JY, Jokitalo E, Sauer N, Scheres B, Nakajima K, Carlsbecker A, Gallagher KL, Helariutta Y (2011) Callose biosynthesis regulates symplastic trafficking during root development. Dev Cell 21 (6):1144–1155. doi:10.1016/j.devcel.2011.10.006

Verna C, Ravichandran SJ, Sawchuk MG, Linh NM, Scarpella E (2019) Coordination of tissue cell polarity by auxin transport and signaling. Elife 8. doi:10.7554/eLife.51061

Yamaguchi YL, Ishida T, Yoshimura M, Imamura Y, Shimaoka C, Sawa S (2017) A Collection of Mutants for CLE-Peptide-Encoding Genes in Arabidopsis Generated by CRISPR/Cas9-Mediated Gene Targeting. Plant Cell Physiol 58 (11):1848–1856. doi:10.1093/pcp/pcx139

Yanagisawa M, Poitout A, Otegui MS (2021) Arabidopsis vascular complexity and connectivity controls PIN-FORMED1 dynamics and lateral vein patterning during embryogenesis. Development. doi:10.1242/dev.197210.

